# Mass-sensitive particle tracking (MSPT) to elucidate the membrane-associated MinDE reaction cycle

**DOI:** 10.1101/2021.04.08.438961

**Authors:** Tamara Heermann, Frederik Steiert, Beatrice Ramm, Nikolas Hundt, Petra Schwille

## Abstract

In spite of their great importance in biology, methods providing access to spontaneous molecular interactions with and on biological membranes have been sparse. So far, it has been consensus that their observation with sufficient sensitivity and time resolution requires the introduction of - predominantly fluorescent-labels to the system. However, the recent advent of mass photometry to quantify mass distributions of unlabelled biomolecules landing on surfaces raised hopes that this approach could be transferred to membranes. Here, we introduce mass-sensitive particle tracking (MSPT), enabling simultaneous label-free tracking and monitoring of molecular masses of single biomolecules diffusing on lipid membranes. We applied this approach to the highly non-linear reaction cycles underlying MinDE protein self-organisation. MSPT allowed us to determine the stoichiometry and turnover of individual membrane-bound MinD/MinDE protein complexes and to quantify their size-dependent diffusion. We found that MinD assembles into complexes larger than the commonly postulated dimer, through lateral interactions of membrane-bound complexes and subunit recruitment from solution. Furthermore, the ATPase-activating protein MinE interconnects MinD into high-molecular-weight heteromeric complexes and affects their subunit turnover and concerted membrane release. This study demonstrates the potential of MSPT to enhance our quantitative understanding of both prokaryotic and eukaryotic membrane-associated biological systems.

## Introduction

The recruitment of proteins to lipid interfaces is crucial for various cell biological processes, such as the regulation of membrane trafficking^1^, mediation of signalling cascades^2^, and the establishment of cell polarity. These membrane-associated reactions often rely on short-lived complexes that coexist in a dynamic equilibrium with their respective cytosolic forms^3^. Transient interactions on the membrane eventually serve as nucleation sites for the assembly of larger, more stable complexes. Additionally, the spatial distribution, stoichiometry and temporal dynamics of membrane-associated complexes are often heterogeneous^4^. This combination of fast dynamics and compositional heterogeneity makes membrane-associated reactions difficult targets for conventional analytical techniques, which determine the composition and follow the dynamics of molecular systems based on ensemble-averaged measures^5^. In recent decades, single-particle tracking (SPT) has revolutionised the analysis of membrane-associated systems by following the dynamics of individual molecules at nanometre precision and millisecond time resolution^6–8^, using fluorophores as labels in combination with highly sensitive microscopy. More recently, scattering-based detection using gold nanoparticles as labels has pushed the spatiotemporal resolution down to the sub-nanometre and microsecond range^9–12^. The single-molecule nature of these approaches has provided detailed mechanistic insight into the dynamics of biomolecular systems^13^. However, as label-based technique, SPT also suffers from label-induced artifacts: large particles, but also small fluorescent tags^14^ may perturb native protein function, and fluorescence-specific phenomena like photo-bleaching and -blinking hinder continuous particle tracking and limit the attainable spatiotemporal resolution^15^. More importantly, although fluorescence-based SPT may provide access to interaction dynamics between tagged molecules through brightness changes, it has proven extremely complicated to extract the molecular composition of the tracked particles. In general, relating the signal of an external marker to the molecular stoichiometry of the labelled particle requires careful characterisation of labelling efficiencies and imaging conditions, which is very often hampered by quenching effects. Especially for multicomponent systems, access to the molecular composition of single particles has been prohibitive for standard biological applications.

Recent advances in interferometric scattering (iSCAT) microscopy^11,16^ made it possible to detect individual biomolecules label-free based on light scattering. The linear relationship between iSCAT contrast and molecular mass of a biomolecule led to the development of mass photometry (MP)^17,18^. This technique allows the determination of size distributions of biomolecules in solution by measuring the individual masses of molecules landing on a glass surface and counting their relative abundances, to determine both molecular identity and composition. For purely shot noise-limited detection of a single molecule landing on a glass surface on top of the static scattering pattern, MP applies a sophisticated background removal strategy optimised for this type of experiment, which is not compatible with the detection of mobile molecules diffusing on lipid membranes. Here, by introducing a new iSCAT image processing and analysis strategy, we enable mass-sensitive particle tracking (MSPT) of single unlabelled biomolecules on a supported lipid bilayer (SLB). We show that the iSCAT signal-to-mass linearity holds true for membrane-associated proteins and that we can relate molecular composition, accessible via mass, to diffusive behaviour. Moreover, we can follow the mass time course along individual trajectories, making it possible to observe the (dis-)assembly of biomolecular complexes in real-time. The approach is fast and provides high particle statistics within minutes, all without need for protein labelling and its caveats.

We showcase the abilities of this method for the detailed analysis of complex biological systems by analysing the membrane-associated reaction cycle of the *Escherichia coli* Min system. This system consists of three proteins - MinC, MinD and MinE - and is essential for the spatiotemporal regulation of the division site in *E. coli*^19^. To perform this task, the ATPase MinD and the ATPase-activating protein MinE oscillate between the cell poles, forming a concentration gradient of the passenger protein MinC. This gradient supposedly enables MinC to inhibit FtsZ polymerisation at the poles and directs Z-ring formation to the mid-cell^20,21^. To this end, the phospholipid bilayer acts as a catalytic interface and membrane interaction of MinD and MinE is mediated by amphipathic helices, i.e. membrane targeting sequences (MTS)^22–25^. The interaction of MinD and MinE with the membrane decreases their diffusion rates and facilitates lateral molecular interactions enabling the system to self-organise^26^. Despite its compositional simplicity the system exhibits surprisingly complex, non-linear dynamics. Its self-organisation can be reconstituted *in vitro*^26^ and the underlying mechanism has been probed by various techniques such as SPT^13^, high-speed atomic force microscopy^27^, nuclear-magnetic resonance^28^, electron microscopy^29^, plasmonic nano sensors^30^ and mutational analysis^31,32^. Despite this intense characterisation, molecular details about MinDE self-assembly, such as the presumed cooperativity in bilayer-attachment of MinD, have remained poorly understood. Thus, the system was set to benefit from the unique ability of MSPT to characterise dynamics as well as molecular composition of membrane-attached protein complexes. Using MSPT, we dissected the membrane-associated MinDE reaction cycle by determining the stoichiometry, turnover, and diffusion of individual membrane-bound MinD/MinDE protein complexes. Our results indicate that MinD, in contrast to the classical model, assembles not only into dimers, but also forms larger oligomers, confirming recent findings^27^. We furthermore show that MinE promotes MinD self-assembly on the lipid bilayer due to its ability to interconnect MinD into so far unresolved higher-order heteromeric complexes. We believe our experiments on the Min system demonstrate that MSPT is a powerful, widely applicable tool for the mechanistic analysis of both pro- and eukaryotic membrane-associated systems.

## Results

### Mass-sensitive dynamic imaging of membrane-associated protein complexes

The major obstacle for detecting single macromolecules with iSCAT microscopy (Fig. 1a) is separating their comparatively small signal from the dominant scattering background. In a standard mass photometry experiment, molecules landing on a glass surface are detected through continuous comparison of an averaged image with the average of its preceding frames as background (Supplementary Fig. 1a)^17,18^. In this image processing approach, a molecule appears as a dark spot at the moment of landing and disappears when it becomes part of the static background^17^. The mass of the particle can then be determined by fitting the detected peak signal with a model point spread function (PSF). However, for a moving molecule, this strategy produces distorted images of the molecule’s PSF. The missing particle density at its previous location produces a bright spot, while the added density at the new position generates a dark spot. The spatial overlap of these patterns causes moving objects to appear as dark fronts carrying bright tails, thus hampering the determination of their mass and location (Supplementary Fig. 1b, Supplementary Movie 1). To address this issue, we employed the temporal median of an image sequence as background estimate^10,33,34^. If during the median period moving molecules only occasionally cross a surface location, the median signal at that location will be a good estimate of the empty background. To retain shot-noise limited detection, which is strongly affected by sample drift, we calculated the pixel-wise median of an image sequence as background estimate for its central frame and moved this median window from frame to frame throughout the video (Supplementary Fig. 1c). Due to the clear separation of static background from moving objects, background corrected movies showed clear, undistorted images of moving PSFs (Fig. 1a, Supplementary Fig. 1c, Supplementary Table 1 and Supplementary Movie 1), a prerequisite that allows mass-sensitive particle tracking (MSPT) of single unlabelled biomolecules diffusing on lipid interfaces.

**Figure 1.**
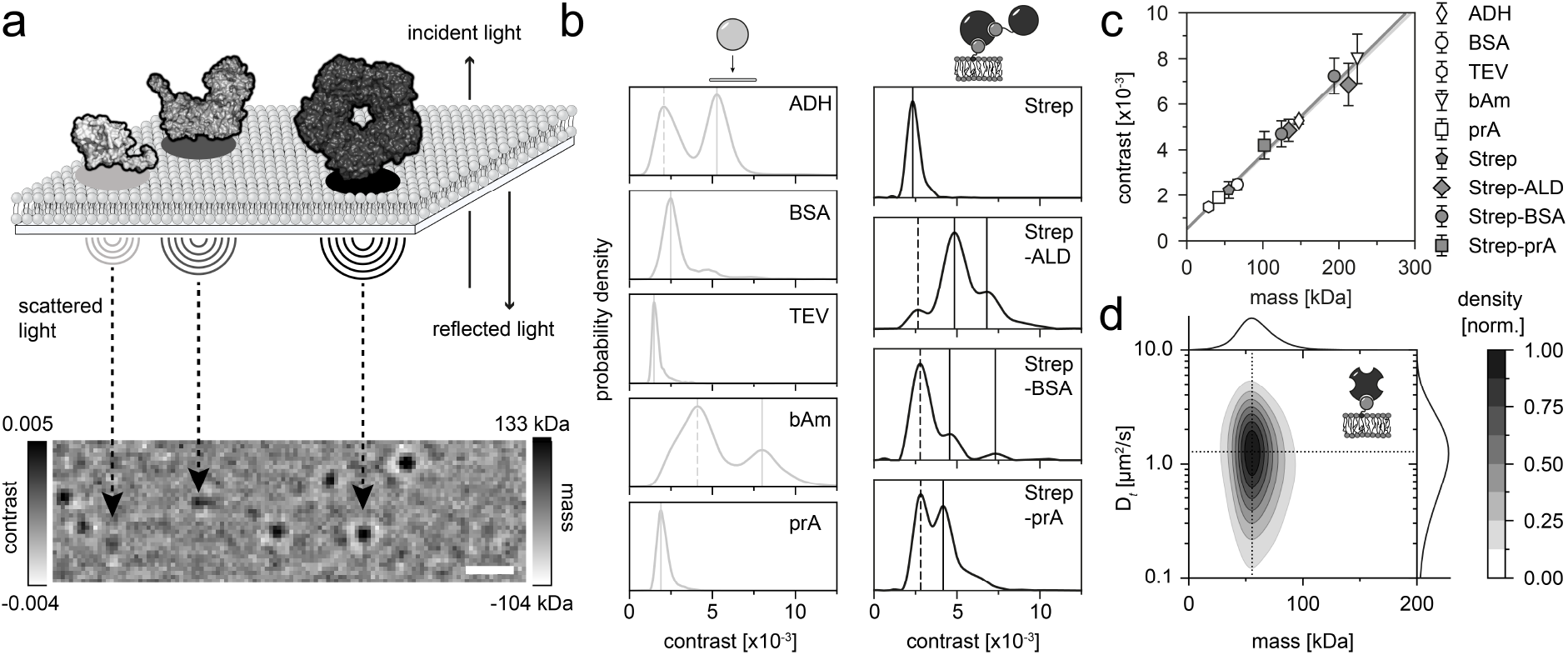
Principle of mass-sensitive particle tracking (MSPT). **(a)** Schematic displaying the interferometric scattering (iSCAT)-based measurement principle of MSPT. In this setup, proteins freely diffuse on a supported lipid bilayer and can be tracked and identified according to the linear relationship between their interferometric scattering contrast and molecular weight. Exemplary structures of three aldolase oligomer states (PDB: 4S1F^40^) are shown in the top panel, and their respective iSCAT images at the bottom. Scale bar: 1 µm. **(b)** Probability density distributions of standard proteins determined using the conventional MP landing assay (left panel) or using MSPT (right panel). All data represent pooled distributions of three independent experiments per condition: alcohol dehydrogenase (ADH; particle number n = 9828), bovine serum albumin (BSA; n = 11,408), TEV protease (TEV; n = 1,705), βamylase (bAm; n = 10,043), protein A (prA; n = 12,720), divalent streptavidin (Strep; n = 16,699 trajectories), divalent streptavidin with biotinylated aldolase (Strep-ALD; n = 16,727 trajectories), divalent streptavidin with biotinylated bovine serum albumin (Strep-BSA; n = 8,842 trajectories) and divalent streptavidin with biotinylated protein A (Strep-prA; n = 22,424 trajectories). Dashed lines mark peaks not considered for mass calibration (left panel). Continuous lines represent oligomer states included in the mass calibration. 2D plots of mass vs. diffusion coefficient for the four proteins measured with MSPT (right panel) are shown in Supplementary Fig. 4. **(c)** Comparison of the contrast-to-mass calibration for MP and MSPT, derived from peak contrasts in panel (b) and their assigned sequence masses (see Supplementary Table 2, 3). Error bars represent the standard error of the peak locations estimated by bootstrapping. **(d)** 2D kernel density estimation of 1.25 nM tetravalent streptavidin bound to biotinylated lipids on a supported lipid bilayer (n = 73,901 trajectories of three independent replicates; particle density: 0.2 µm^−2^). Marginal probability distributions of the molecular mass and the diffusion coefficient are presented at the top and right, respectively.

First, we set out to assess the quality of mass determination of mobile molecules compared to landing particles in conventional mass photometry. To this end, we used a bilayer supplemented with biotinylated lipids and attached a set of biotinylated standard proteins with known mass via divalent streptavidin^35^. This system has several advantages, such as the ability to cover different protein size regimes, standardised membrane binding, and the added benefit of simplified complex stoichiometries by using divalent streptavidin. For each molecule, we determined its median contrast throughout its trajectory. Analogous to conventional mass photometry (Fig. 1b, left column), histograms of the contrasts of all molecules determined with MSPT revealed the particle size distribution (Fig. 1b, right column, Supplementary Movie 2). For our standard proteins, iSCAT contrast as a function of mass exhibited the expected linear relationship^17^. The calibration line obtained from standards diffusing on SLBs was in fact indistinguishable from the one determined with molecules landing on glass (Fig. 1c). This result suggests that mass calibrations performed with landing assays can be transferred to particles diffusing on membranes. However, this strictly needs to be verified for any new lipid/buffer combination, protein system and imaging condition.

Besides mass determination, MSPT also enables the analysis of the diffusive behaviour of membrane-bound molecules. For this purpose, it is important to choose the median window size for background estimation such that particles travel sufficient distances during the median period. Hence, we first systematically tested the minimum median window sizes required to extract the correct diffusion coefficients at particle diffusion speeds expected for our SLB system. We generated artificial movies of randomly diffusing particles at varying speeds and compared the input diffusion coefficients with diffusion coefficients extracted using different median window sizes and extraction methods (see methods, Supplementary Figures 2 and 3 as well as Supplementary Discussion for details). Based on the results of our simulation, we chose a jump-distance analysis^36^ to extract diffusion coefficients from our experimental videos. As an experimental verification of the diffusion coefficients obtained in this manner, we again made use of streptavidin attached to a membrane via biotinylated lipids. In line with literature values ranging from 0.8 to 2.0 µm^2^ s^−1 37–39^, the lateral diffusion coefficient of streptavidin was found to be 1.3 ± 0.1 µm^2^ s^−1^ (Fig. 1d). Similar diffusion coefficients were obtained for all standard proteins attached via divalent streptavidin (Supplementary Fig. 4). To highlight a unique advantage of MSPT, we plotted the diffusion coefficient versus the respective molecular mass obtained from individual trajectories, enabling an unprecedented direct connection of these two parameters. As displayed in Figure 1d, the unimodal distribution of membrane-bound streptavidin indicates a distinct population of tetramers undergoing Brownian motion. Having validated the method with membrane-attached streptavidin, we thought the method offers the potential of detailed insight into the dynamics of molecular interactions within more complex membrane-bound systems. To further explore the method’s capabilities, we turned to the membrane-associated *E. coli* Min system.

### Cooperative Membrane-Catalysed Association Dynamics of MinD

The MinDE system is known for its ability to self-organise into mesoscopic protein patterns on lipid membranes^26^. To generate these patterns, MinD and MinE are generally assumed to undergo a canonical membrane binding-unbinding cycle displayed in Figure 2a. In brief, upon ATP-complexation cytosolic MinD dimerises and localises to the membrane interface^23^. After homodimeric MinE binds to MinD, nucleotide hydrolysis is stimulated and MinD dissociates from the lipid bilayer to return to its monomeric state^41^. Despite this established model, it has remained rather enigmatic how ATP-dependent dimerisation and the resulting increase in MinD membrane affinity alone can confer the non-linear attachment required for pattern formation. In recent years, it has been proposed that the presence of higher MTS valences and thus higher-order oligomer structures might contribute to the local self-enhancement of MinD at the membrane^27,31,42^. Due to the fast diffusion and attachment/detachment dynamics, however, it has remained challenging to provide convincing evidence of their existence.

**Figure 2.**
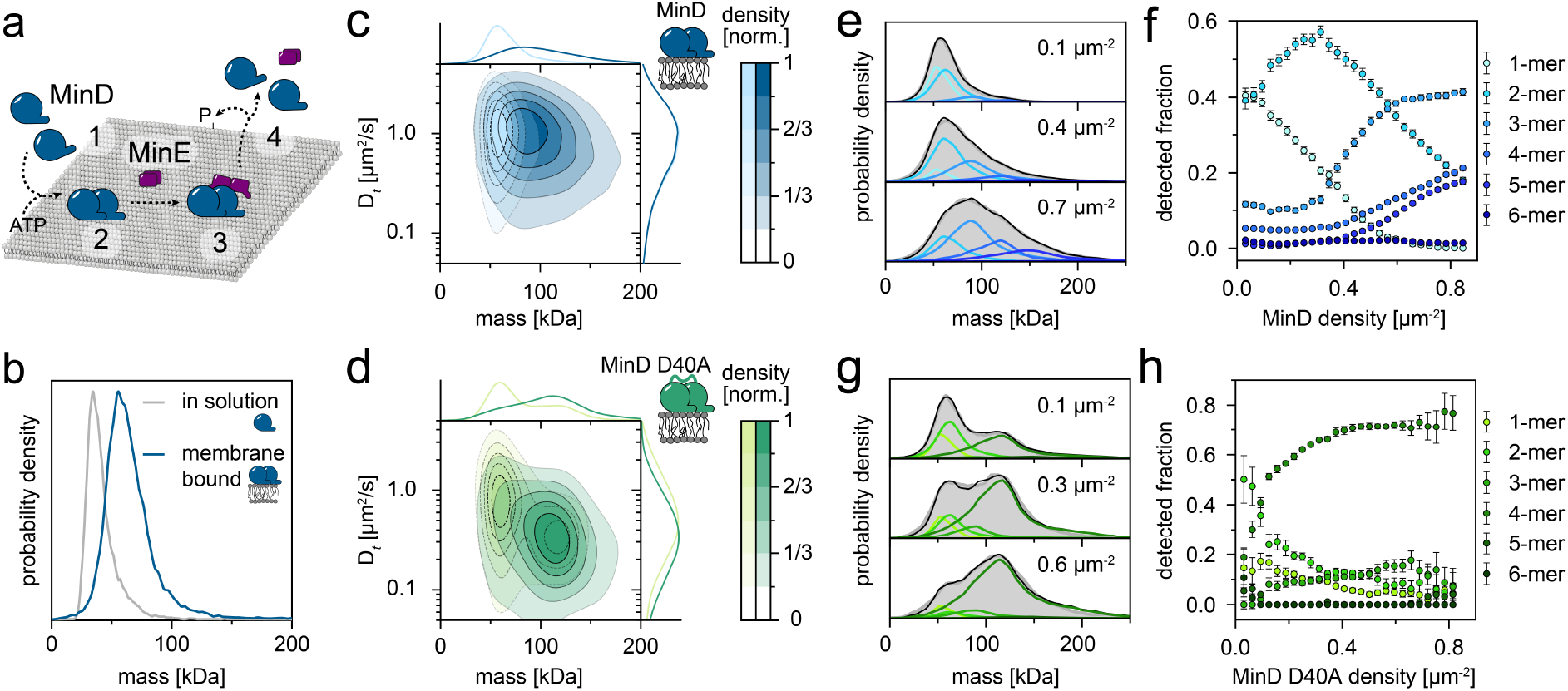
Lateral MinD–MinD interactions lead to self-assembly into large homo-oligomers. **(a)** Schematic of the canonical membrane binding-unbinding cycle of MinDE. Upon ATP-complexation, MinD dimerises (1) and attaches to the membrane interface (2). In the event of MinE binding (3), MinE stimulates the intrinsic ability of MinD to hydrolyse ATP, which leads to the release of MinD from the membrane in its monomeric form (4). **(b)** MinD mass distribution in solution (grey line; n = 16,101 trajectories) and upon attachment to the supported lipid bilayer (blue line; n = 13,917 trajectories). For solution experiments, 175 nM MinD with 0.5 µM ATP were measured in the conventional MP landing assay. The membrane mass distribution of MinD was determined using MSPT at a particle density of 0.03 µm^−2^. **(c)** 2D kernel density estimation of membrane-attached MinD and **(d)** MinD D40A at particle densities of 0.1 µm^−2^ (light blue, n = 117,086 trajectories) and 0.8 µm^−2^ (dark blue, n = 152,685 trajectories) and 0.1 µm^−2^ (light green, n = 7,831 trajectories) and 0.8 µm^−2^ (dark green, n = 3,150 trajectories), respectively. Marginal probability distributions of both molecular mass and diffusion coefficient are presented at the top and right, respectively. **(e)** Representative mass distributions (grey) of MinD and MinD D40A **(g)** and estimation (black line, coloured lines highlight underlying components) of its six components for three different particle densities. MinD: 0.1 µm^−2^ – light blue, 0.4 µm^−2^ – blue, 0.7 µm^−2^ – dark blue; MinD D40A: 0.1 µm^−2^ – light green, 0.3 µm^−2^ – green, 0.6 µm^−2^ – dark green. **(f**,**h)** Relative oligomer abundance as a function of particle density. Error bars are the standard deviation of fitting results from three data subsets. The oligomer analysis is based on a total of n = 1,102,940 trajectories for MinD and n = 194,545 for MinD D40A.

Taking advantage of the ability to monitor the mass of membrane-attached and diffusing protein complexes, we thus applied our MSPT approach to investigate potential oligomeric species formed by the MinDE system. We started by reconstituting the membrane recruitment of MinD and compared the mass distribution of MinD in solution, measured in an MP landing assay, with the mass distribution of MinD on a lipid bilayer (Fig. 2b, Supplementary Fig. 5). In the presence of ATP, MinD monomers were detected as the predominant species in solution. In contrast, the major species on the membrane was a MinD dimer already at sparse densities corresponding to one particle per frame in the field of view (31.9 µm^2^). However, it should be noted that at the chosen imaging conditions for MSPT, the signal-to-noise ratio of monomers (33 kDa) was too low for their quantitative detection (Supplementary Figures 9 and 10). Hence, an adequate estimate of their membrane abundance cannot be stated.

One of the major advantages of iSCAT-based imaging is that it provides a direct estimate of the molecular density on a bilayer from the number of detected particles, as compared to single-molecule fluorescence where factors like labelling efficiency and photo bleaching have to be taken into account. Accordingly, we could classify video sections of membrane-bound MinD into conditions of different particle densities and observed the resulting change in the oligomeric distribution (Fig. 2c, Supplementary Fig. 6, 7 and Supplementary Movie 3). While MinD was mainly present in the dimer state at low particle densities (0.1 µm^−2^), the MinD population shifted towards a broad distribution with higher-order complexes on crowded bilayers (0.8 µm^− 2^). This data directly demonstrates that MinD indeed assembles into complexes larger than a dimer as previously suggested^27,31^. To further investigate the structural determinants for higher-order MinD assemblies, we performed the same MSPT experiment with a MinD mutant (D40A) that is reported to predominantly reside in the dimeric state due to its impaired ability to hydrolyse ATP^43^. According to the prevailing mechanism, it should not be possible for a locked dimer mutant to participate in MinD self-assembly, due to its inability to switch between the monomeric and dimeric state^23,41,44^. Nevertheless, we found a distinct population of MinD D40A tetramers in our contour plot (Fig. 2d, Supplementary Fig. 8), suggesting that MinD is capable of forming higher-order oligomers using an interface distinct from the canonical dimerisation site.

Another striking aspect of the D40A mutant was its clear separation into two distinct populations (dimer and tetramer), whereas the distribution of the WT protein appeared unresolved. This result suggested that the WT was able to also recruit MinD monomers forming trimeric species as intermediates, which were not as abundant for the D40A mutant. In order to provide a more quantitative measure for the comparison between WT and D40A mutant, we deconvolved their mass distributions into the underlying components to determine the relative abundance of each species as a function of particle density on the bilayer. To estimate the shapes of the individual components, we used our video simulation routine and determined separate mass distributions for particles representing only one type of MinD oligomer at a range of particle densities (Supplementary Figures 9, 10 and Supplementary Discussion). Next, we fit the mass distributions of MinD WT and D40A using a linear combination of simulated distributions from six components corresponding to monomers – hexamers and extracted the relative abundance of each oligomer as a function of particle density (Fig. 2e, g, Supplementary Fig. 11). These graphs show a sequential appearance of increasingly larger oligomers of MinD for higher molecule densities (Fig. 2f). Compared to the WT, the D40A mutant had a higher tendency to populate stoichiometries with even numbers of subunits and to transform its dimer state into the tetramer state, likely due to the increased stability of the dimer (Fig. 2h).

Furthermore, MSPT confers the unique possibility to determine oligomer-specific lateral diffusion coefficients. This knowledge can be used to deduce structural information about the observed molecules when considering the theory of Evans-Sackmann^45^, which postulates the relation of an object’s membrane inclusion size with its respective diffusion coefficient. Assuming a similar membrane viscosity as for pure DOPC membranes^46^, we can thus estimate that a MinD D40A dimer with a diffusion coefficient of 0.85 µm^2^/s has an inclusion size of 5 nm, and a tetramer with 0.34 µm^2^/s an inclusion size of 9 nm (Supplementary Fig. 13). These estimates imply that all monomers^43^ of the tetramer are able to insert their MTS into the bilayer, suggesting certain geometrical constraints for subunit orientation.

### Time-resolved mass analysis of single MinD trajectories

Aside from measuring a particle’s location frame by frame, MSPT also allows to determine its respective mass in a time-resolved fashion, thus enabling the detection of attachment and detachment events along the trajectory of a single particle (Fig. 3a, Supplementary Fig. 14). For MinD, the minimal expected mass increment (33 kDa) for monomer-wise turnover was close to the measurement uncertainty (28 kDa standard deviation) of the mass for a single frame. To minimise user bias, we employed a step-finding algorithm that locates mass change points along trajectories based on statistical criteria^47^.

**Figure 3.**
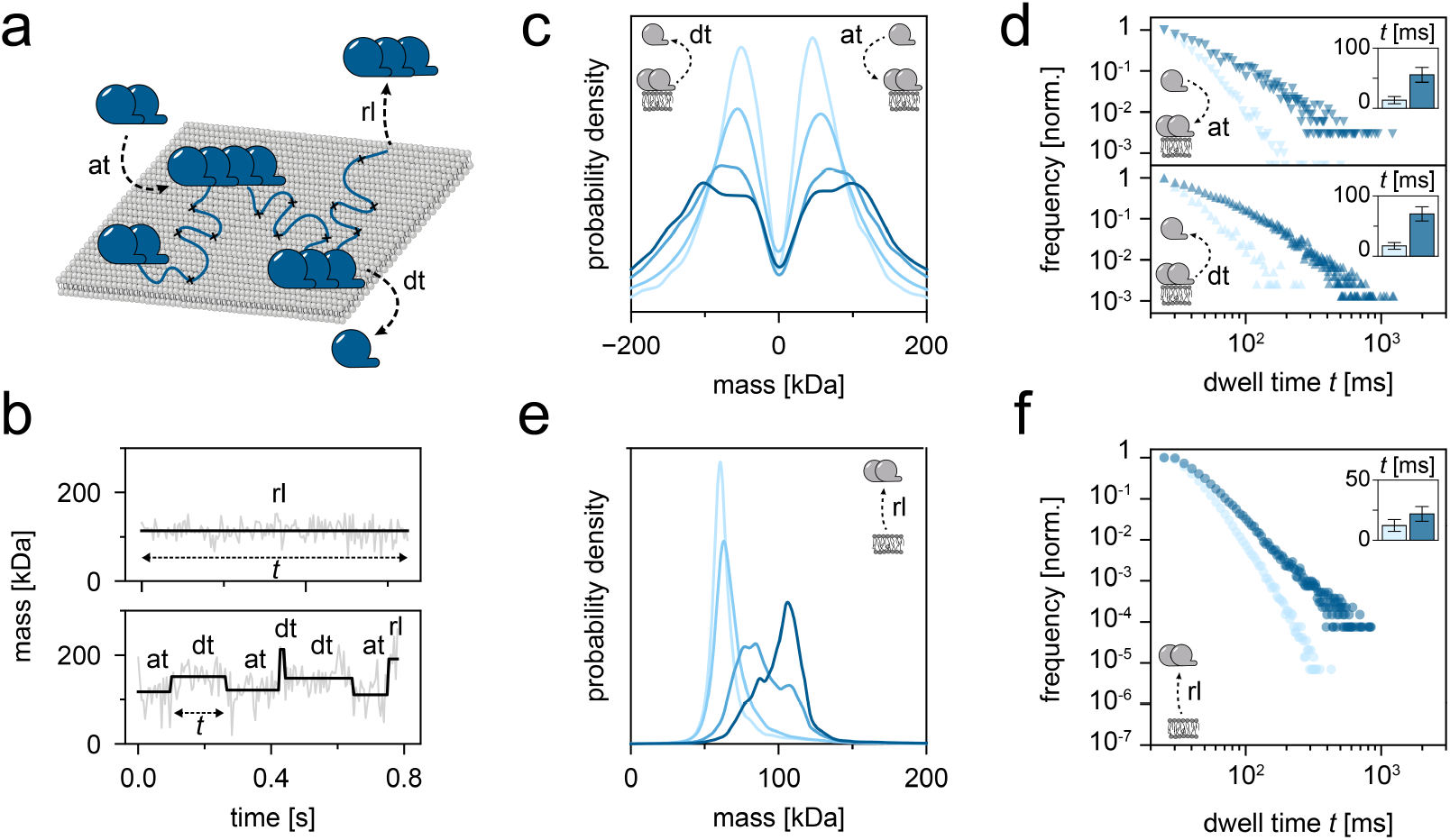
Subunit (dis-)assembly of MinD particles diffusing on membranes. **(a)** Schematic representation of the time-resolved mass analysis of single MinD trajectories, which reveals attachment (at) and detachment (dt) events along the trajectory as well as a particle’s full membrane release (rl) at the end of its trajectory. **(b)** Representative mass time traces of MinD trajectories (grey line) and corresponding step fits (black line) determined by a step-finding algorithm that locates mass change points within a trajectory. **(c)** Mass step size distribution derived from step fits as depicted in (b) revealing MinD subunit sizes for attachment (at) and detachment (dt) events at particle densities: 0.1 µm^−2^ – pale blue (n = 20,796 plateaus), 0.3 µm^−2^ – light blue (n = 26,177 plateaus), 0.6 µm^−2^ –blue (n = 25,864 plateaus), 0.8 µm^−2^ –dark blue (n = 7,506 plateaus). **(d)** Dwell time plots for MinD particles before attachment (at) events (top plot), representing the lengths of mass plateaus preceding a mass increase, and before detachment (dt) events (bottom plot), representing the lengths of mass plateaus preceding a mass decrease. Dwell times are shown for the MinD dimer (66 kDa, light blue line) and the MinD tetramer state (132 kDa, dark blue line). Plateau numbers for (at): dimer - n = 23,782; tetramer - n = 5,088; for (dt): dimer - n = 3,143; tetramer - n = 10,406. Insets: Mean dwell times of the two oligomer states. Error bars represent the standard deviation of the dwell times estimated by bootstrapping. **(e)** MinD mass distribution for membrane release (rl) events. MinD particle densities: 0.1 µm^−2^ – pale blue (n = 117,086 plateaus), 0.3 µm^−2^ – light blue (n = 169,957 plateaus), 0.6 µm^−2^ – blue (n = 284,916 plateaus), 0.8 µm^−2^ – dark blue (n = 120,654 plateaus). **(f)** Plot of the dwell times before membrane release (rl) for the MinD dimer and tetramer state. Plateau numbers: dimer - n = 562,011; tetramer - n = 73,037. Inset: Mean dwell times of the two oligomer states. Error bars represent the standard error of dwell times estimated by bootstrapping.

Figure 3b displays examples of mass time courses for individual particles and the mass steps detected by the algorithm. Note that in a majority of cases, MinD complexes retained their size throughout their entire trajectory (Fig. 3b, upper panel). However, in ~10% of trajectories, the mass of a tracked particle changed during its trajectory (Figure 3b, bottom panel), suggesting the attachment or detachment of MinD subunits to and from the membrane-bound complex. By analysing the sizes of mass steps along all trajectories, it was possible to obtain a distribution of subunit sizes attaching and detaching from membrane-bound MinD complexes (Fig. 3c), which appeared to mostly (dis-)assemble in one- or two-subunit increments at low particle densities. However, for higher MinD particle densities, when larger oligomers had accumulated, these complexes often turned-over greater subunits, as indicated by the shift of the distribution towards higher mass steps^27^ (Fig. 3c, dark blue profile). Moreover, the combined information of a mass plateau level and its dwell time as annotated in Fig. 3b could be used to extract the subunit turnover rates of each oligomer species (Fig. 3d). Here, the dwell time preceding a mass increase could be used to deduce subunit attachment rates (Fig. 3d – upper panel), whereas the dwell time followed by a mass decrease provided an estimate of detachment rates (Fig. 3d – lower panel). The resulting lifetime plots suggested that membrane-attached dimers had a faster subunit turnover than tetramers, implying higher stability of these larger complexes.

Plateaus at the end of trajectories (Fig. 3b, bottom panel, last plateau), and trajectories without any mass change at all (Fig. 3b, top panel), could be used to identify the molecular weight of particles completely released (rl) from the membrane (Fig. 3e). Notably, this time-resolved mass analysis of the individual trajectories significantly improved the mass resolution, as compared to the median-based particle mass estimates used in Figure 2. Hence, MinD dimers, trimers and tetramers were now fully resolved as separate peaks (Fig. 3e). Accordingly, one could now recognize that the major species released from the membrane was a dimer at low particle densities and a tetramer at high particle densities. The corresponding dwell time plot showed that tetramers stayed associated to the bilayer significantly longer than dimers, in line with a higher avidity in membrane binding conferred by additional MTS (Fig. 3f). For comparison, the dimer-arrested mutant MinD D40A almost exclusively dissociated as dimers or tetramers (Supplementary Fig. 15, 16). Taken together, our detailed trajectory analysis confirms our previous observations suggesting that MinD WT assembles into species of higher order with an intermediate trimer state. This indicates that subunits bind at a location different from the canonical dimerisation site.

### MinE-induced formation of large heteromeric complexes

In the past decade, several different mechanistic models have been proposed to explain the role of the ATPase-activating protein MinE for MinDE detachment dynamics^13,48,49^. Some of these models are based on the cooperation of both MinE and MinD dimer to prompt membrane-release through MinD ATPase activity stimulation^44,48^. However, this effect alone cannot explain the recently observed cooperative membrane-detachment of MinDE filaments^27^. To address this issue, we used MSPT to determine the stoichiometry of the membrane-bound MinDE complex and followed its membrane dynamics on a molecular level.

In accordance with previous structural studies that suggest a conformational switch of MinE allowing MinD binding only upon the encounter of membrane-bound MinD^48,50^, we found no indication for MinDE interaction in solution (Supplementary Fig. 17). In the presence of a supported lipid bilayer, however, the MinDE complex existed predominantly in a stable double-dimeric state (Fig. 4a – light pink, Supplementary Fig. 18). Furthermore, if the MinDE complex encountered more proteins on a crowded bilayer, MinE promoted the interconnection into very large heteromeric MinDE complexes, a behaviour unexpected considering the common models (Fig. 4a – magenta). One possible explanation for this behaviour is the ability of a MinE dimer to symmetrically bind to both sides of a MinD dimer, thus effectively acting as a bridge between MinD assemblies^49^. Accordingly, our time-resolved mass step analysis revealed that during subunit turnover on membrane-bound particles, predominantly dimeric and tetrameric subunits attached (at) and detached (dt) at high particle densities of 0.6 and 0.8 µm^−2^ (Fig. 4b, Supplementary Fig. 19). This effect required a critical minimum density on the bilayer, since MinDE complexes in sparsely populated environments (0.1 and 0.3 µm^−2^) mainly exhibited conversion in their minimum subunit increments. Hence, their final oligomeric state during membrane release (rl) resembled a MinDE dimer (Fig. 4c, light pink). Strikingly, on bilayers with high protein density, MinDE complex sizes released from the membrane increased beyond the sizes observed for MinD alone and reached masses of >350 kDa (Fig. 4c, purple).

**Figure 4.**
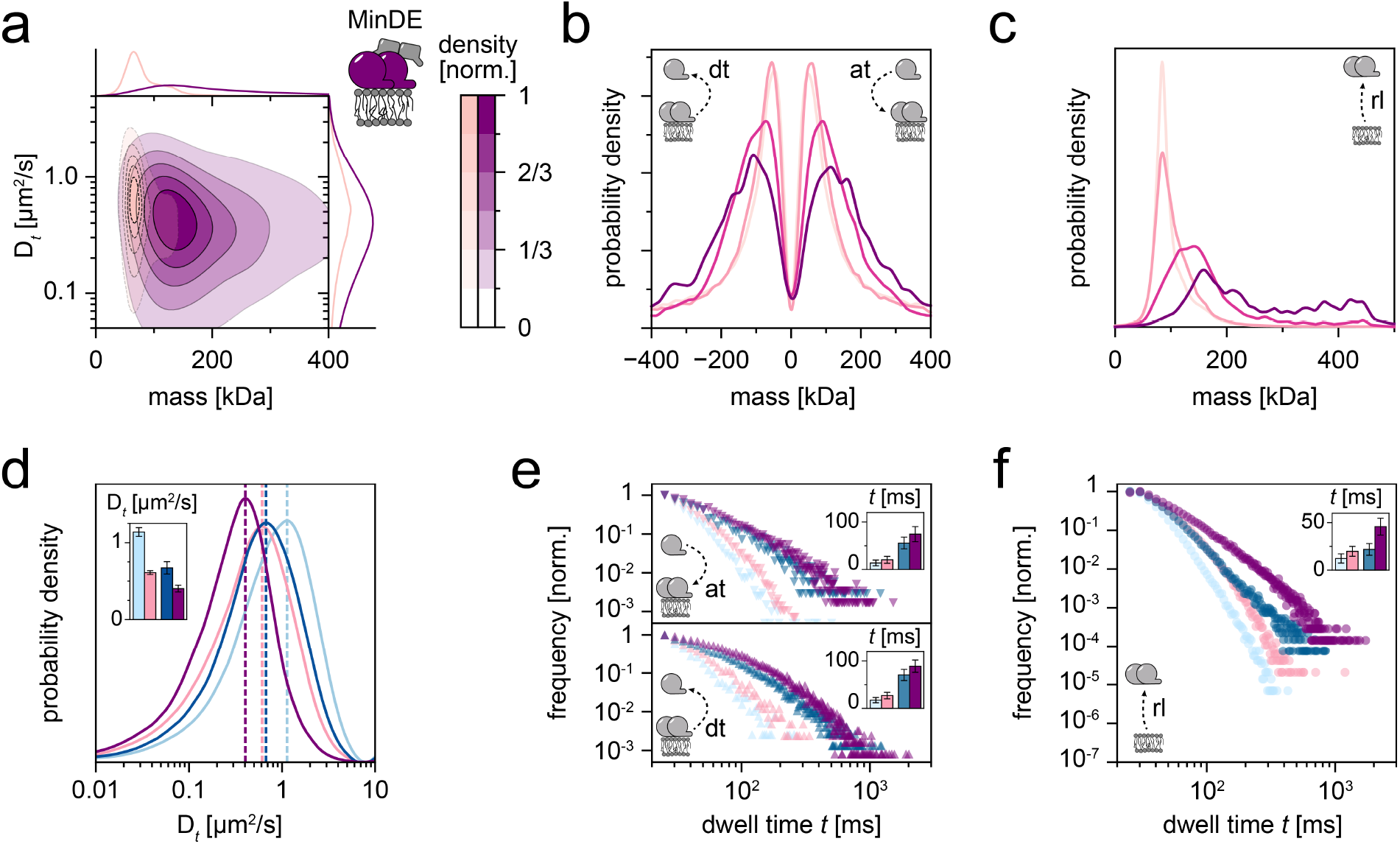
MinE interconnects MinD oligomers into large complexes with a prolonged membrane dwell time. **(a)** 2D kernel density estimation of membrane-attached MinDE complexes at particle densities of 0.1 µm^−2^ (light pink, n = 200,436 trajectories) and 0.8 µm^−2^ (purple, n = 30,082 trajectories). Marginal probability distributions of both molecular mass and diffusion coefficient are presented at the top and right, respectively. **(b)** Mass step size distribution revealing MinDE subunit turnover (at and dt events) on membrane-bound particles at particle densities of 0.1 µm^−2^ – pale pink (n = 105,438 plateaus), 0.3 µm^−2^ – light pink (n = 73,471 plateaus), 0.6 µm^−2^ – pink (n = 9,247 plateaus), 0.8 µm^−2^ – dark purple (n = 4,040 plateaus). **(c)** MinDE mass distribution for membrane release (rl) at MinDE membrane particle densities of: 0.1 µm^−2^ – pale blue (n = 200,436 plateaus), 0.3 µm^−2^ – light blue (n = 158,660 plateaus), 0.6 µm^−2^ – blue (n = 35,501 plateaus), 0.8 µm^−2^ – dark blue (n = 26,708 plateaus). **(d)** Analysis of oligomer specific diffusion coefficients for MinD (blue lines) and MinDE complexes (pink lines). light blue/pink – dimer (MinD/MinDE n = 439,568/206,422 trajectories); dark blue/purple – tetramer dimer (MinD/MinDE n = 118,922/47,136 trajectories). Inset: Mean diffusion coefficients of each oligomer state. Error bars represent the standard deviation of the dwell times estimated by bootstrapping. **(e)** Dwell time plots for MinD and MinDE attachment (at) events (top plot) as well as for MinD and MinDE detachment (dt) events (bottom plot). Dwell times are shown for the MinD dimer (light blue line) and MinD tetramer (dark blue line) as well as for their respective MinDE versions (hetero-dimer state – light pink, heterotetramer state – purple). Plateau numbers for (at): MinD dimer – n = 23,782, MinDE dimer – n = 37,278, MinD tetramer – n = 5,088, MinDE tetramer – n = 11,698; (dt): MinD dimer – n = 3,143, MinDE dimer – n = 3,974, MinD tetramer – n = 10,406, MinDE tetramer – n = 22,501. Insets: Mean dwell times of the two oligomer states. Error bars represent the standard deviation of the dwell times estimated by bootstrapping. **(f)** Plot of the dwell times before membrane release (rl) for the MinDE dimer and tetramer state and the respective MinD versions for comparison. Plateau numbers: MinD dimer – n = 562,011, MinDE dimer – n = 277,782, MinD tetramer – n = 73,037, MinDE tetramer – n = 60,114. Inset: Mean dwell times of the two oligomer states. Error bars represent the standard error of the dwell times estimated by bootstrapping.

Based on our mass dwell time analysis, we found that the presence of MinE generally reduced the diffusion coefficients (Fig. 4d) of MinDE oligomers and slowed down turnover rates (Fig. 4e, pink), when compared to their respective MinD versions (Fig. 4e, blue). In addition, MinDE complexes were found to reside significantly longer on the membrane before full release (Fig. 4f). This suggests that MinE can effectively stabilise membrane-bound MinD when present in equimolar amounts as previously suggested^13,49^.

## Conclusions

In this work, we presented a simple and versatile single-molecule-based method for the determination of membrane-associated oligomer distributions, subunit turnover and mass-resolved residence times in the context of the *E. coli* Min system. Based on our results, we propose the extension of the established membrane binding-unbinding models for MinDE self-organisation^44,48^ (Fig. 5). Our data support the original model regarding nucleotide exchange from ADP to ATP triggering membrane-dependent dimerisation of MinD, thus coupling the process to energy dissipation. On the membrane, we find that lateral MinD interactions as well as recruitment of MinD subunits from solution lead to the formation of a dynamic mixture of MinD oligomeric states that assemble through attachment and detachment of subunits at a location different from the canonical dimerisation site^31^, a behaviour unexpected by common models. We assume that the ability of MinD to assemble into these complexes is generally required for its local self-accumulation and in combination with the initial nucleotide-dependent membrane recruitment explains the observed attachment cooperativity during MinDE self-organisation. At elevated membrane densities, a small percentage of MinD self-assembles into tetramers, which either fall apart or interact with MinE. In contrast to the previously assumed cooperation of the MinE and MinD dimer to prompt membrane-release^44,48^, we found that MinE promotes the interconnection^49^ of heteromeric MinDE complexes. We assume that this behaviour is based on the ability of two MinE dimers to symmetrically bind to both sides of a MinD dimer, thus effectively acting as a bridge between MinD assemblies^49^. The existence of multivalent membrane-bound structures would likewise explain the prolonged residence times of MinDE complexes compared to their respective MinD oligomer variants. Notably, opposed to the previously assumed monomer detachment^51^, we found MinDE to detach from the membrane interface in complexes with sizes beyond 350 kDa, which could correspond to eight MinDE subunits. This corroborates that MinE could induce nucleotide-conversion of MinD subunits and thereby weaken the overall membrane avidity of the MinDE complex prior to its full dissociation from the membrane interface^27^. Upon membrane release, larger complexes might remain temporarily stable, stay in the vicinity of the bilayer, and can potentially rebind close to their dissociation spot, once MinD subunits have exchanged their nucleotide. An indication of such behaviour is the gradual shift of MinD/MinDE mass distributions, for both membrane recruitment and release events, towards higher masses over the course of a video (Supplementary Fig. 20). To conclude, we believe that MinDE self-organisation arises from an interplay of cooperative MinD membrane attachment into higher-order oligomers, anisotropy of the local MinD concentration through quick re-binding of oligomers to the membrane, and ATP-dependent membrane release of MinD assemblies coordinated by the ATPase-stimulating activity of MinE.

**Figure 5.**
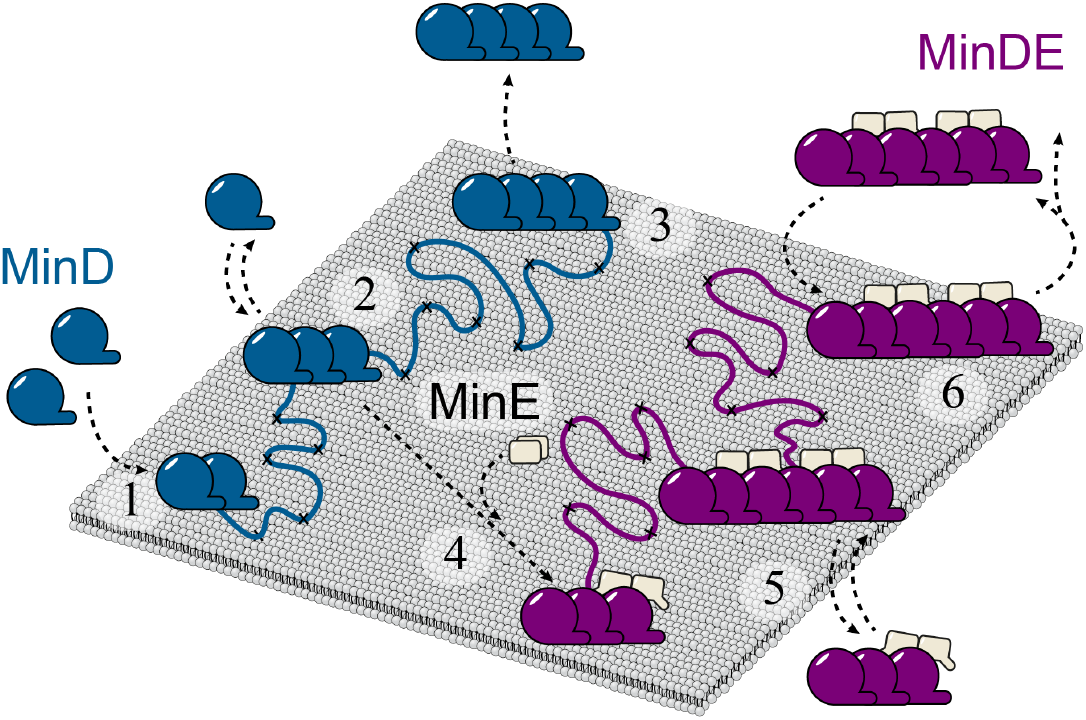
Schematic of the proposed membrane-associated MinDE reaction cycle. **(1)** Upon nucleotide exchange from ADP to ATP, membrane-dependent dimerisation of MinD is triggered. **(2)** On the membrane, lateral MinD interactions and recruitment of MinD subunits from solution lead to MinD higher-order structures that assemble through attachment of subunits at a location different from the canonical dimerisation site. MinD assemblies then either dissociate from the membrane **(3)**, or encounter MinE **(4)**. MinE promotes the interconnection of very large heteromeric MinDE complexes that, due to their multivalent MTS structure, reside significantly longer on the membrane interface **(5). (6)** However, MinE also induces nucleotide-conversion of MinD subunits, thereby weakening the overall membrane avidity of the MinDE complex, prior to its full release from the membrane interface in complexes >350 kDa.

The mechanistic dissection of the MinDE membrane-cycle described in this study constitutes a detailed technical demonstration of the capabilities of mass-sensitive particle tracking (MSPT). By combining the advantages of mass photometry^17,18^, where biomolecular complex stoichiometries are determined from the mass of individual particles, with single-particle tracking, MSPT provides invaluable insights into the complex dynamics of biomolecules on lipid membranes. When exploring the size range for quantitative characterisation of transient membrane interactions and oligomer stoichiometries, we found that >50 kDa proteins on supported lipid bilayers could be reliably resolved with our system at relatively low particle densities (see Supplementary Discussion). Although the application to even smaller proteins will require improved methodology with regard to detection and data processing, the ability of label-free detection of membrane complexes provides powerful advantages over conventional fluorescence-based single-particle tracking approaches: 1) The possibility to simultaneously determine particle densities, i.e. local concentrations on the bilayer; 2) The correlation of mass and diffusion coefficient, which, in combination with a derived membrane inclusion size, provides information about the stoichiometry and arrangement of a membrane-bound complex; 3) The analysis of time-resolved subunit turnover and its kinetics by analysing mass changes along a particle’s trajectory; 4) Extended observation times due to the absence of photo-bleaching, thus enabling the collection of high particle statistics in a short period of time. Ultimately, we believe that mass-sensitive particle tracking will make a strong contribution to the quantitative understanding of both prokaryotic and eukaryotic membrane-associated biological systems.

## Methods

### ADP/ATP stock solution

Both ADP and ATP stocks were prepared from their respective salt hydrates (#A2754 and #A2383, Sigma Aldrich, St. Louis, USA), supplemented with an equal molar amount of MgCl_2_ and adjusted to pH 7.5 with 1 M Tris-HCl. Final nucleotide concentration was spectroscopically determined (*λ* = 259 nm; V-650, Jasco, Pfungstadt, Germany) using an extinction coefficient of 15,400 M^−1^ cm^1^.

### Protein Purification and Modification

Purifications of MinD, MinD D40A and MinE were performed as previously described ^52^ and all used plasmid constructs are indicated in Supplementary Table 4. Divalent streptavidin was assembled as outlined in Howarth *et al*. ^53^, both pET21a-Streptavidin-Alive (Addgene plasmid #20860) and pET21a-Streptavidin-Dead (Addgene plasmid #20859) were a gift from Alice Ting^53^. Biotin-modification of aldolase (#28403842, Cytiva, Marlborough, USA) with EZ-Link™ Maleimide-PEG2-Biotin (#A39261, Thermo Fisher Scientific, Waltham, USA) was achieved through incubation at RT for 1 h and subsequent size-exclusion chromatography on a 16/600 Superdex 200 pg column (GE Healthcare, Pittsburgh, USA), equilibrated in storage buffer (50 mM HEPES pH 7.25, 150 mM KCl, 0.1 mM EDTA, 10% glycerol, 0.4 mM TCEP), using an Äkta Pure chromatography system (GE Healthcare, Pittsburgh, USA). LC-MS and SDS-PAGE was performed to assess purity and integrity of all purified or modified proteins. A customized Bradford assay (#5000006, Bio-Rad Protein Assay; Bio-Rad Laboratories Inc., Hercules, USA) was used to determine protein concentrations and single-use aliquots were flash frozen in liquid nitrogen and stored at −80 °C.

### Small Unilamellar Vesicles (SUVs) and Supported Lipid Bilayer (SLB) Formation

For the formation of small unilamellar vesicles (SUV), dioleoyl-sn-glycero-3-phosphocholine (DOPC; #850375, Avanti Polar Lipids, Alabaster, USA), dioleoyl-sn-glycero-3-phosphoglycerol (DOPG; #840475, Avanti Polar Lipids, Alabaster, USA) and 1,2-dioleoyl-sn-glycero-3-phosphoethanolamine-N-cap biotinyl (18:1 Biotinyl Cap PE; #870273, Avanti Polar Lipids, Alabaster, USA) were dissolved in chloroform (Sigma Aldrich, St. Louis, USA) and mixed in a ratio of 70 mol % DOPC to 30 mol % DOPG or 70 mol % DOPC with 29.99 mol % DOPG and 0.01 mol % Biotinyl Cap PE. After solvent evaporation through nitrogen, residual chloroform was removed for 1 h in a vacuum desiccator. Lipid film hydration was achieved with Min buffer (25 mM Tris/HCl pH 7.5, 150 mM KCl, 5 mM MgCl_2_) and SUVs were formed through consecutive freeze-thaw cycles (8-10) using liquid nitrogen and a 90 °C water bath. For monodisperse vesicle distribution, lipid mixtures were extruded across a Whatman nucleopore membrane (#110603, GE Healthcare, Chicago, USA) with a pore size of 50 nm for 37 passes.

SLBs were formed by fusion of SUVs on cleaned glass cover slides (Paul Marienfeld GmbH & Co. KG, Lauda-Königshofen, Germany) that were assembled into a flow chamber through double-sided sticky tape (Scotch, Conrad Electronic SE, Germany). Prior to their assembly, cover slides (#0102242, #1.5, 24 × 60 mm; #0102062, #1.5, 24 × 24 mm) were cleaned by sequential sonication in Milli-Q water, isopropanol, and Milli-Q water (15 min each), and subsequently dried under a nitrogen stream. Slides were then activated for 30 s (30% power, 0.3 mbar) in a Zepto plasma cleaner (Diener electronic GmbH, Ebhausen, Germany) using oxygen as the process gas. After flow chamber assembly, SUVs were added to each reaction chamber at a final concentration of 0.4 mg/mL in Min buffer with additional 2 mM CaCl_2_ to promote vesicle rupture. Unfused SUVs were removed through subsequent washing with Min buffer.

### Mass Calibration Curves and Diffusion Control

To convert interferometric scattering contrast into the respective protein mass, the contrast of a set of mass standards was measured for each experimental setup – MP landing assays and MSPT on the lipid interface. In line with Young *et al*. ^17^, mass calibration for landing assays was performed in flow chambers filled with filtered (sterile syringe filters 0.45 *μ*m cellulose acetate membrane, VWR International, Radnor, USA) SLB buffer (25 mM Tris/HCl pH 7.5, 150 mM KCl). As mass standards, either 50 nM TEV protease (MPIB Core Facility), 50 nM Pierce™ recombinant protein A (#21184, Thermo Fisher Scientific, Waltham, USA) 50 nM bovine serum albumin (#A4612, Sigma Aldrich, St. Louis, USA), 20 nM alcohol dehydrogenase (#A8656, Sigma Aldrich, St. Louis, USA), or 20 nM β-amylase (#A8781, Sigma Aldrich, St. Louis, USA) were injected and landing events recorded. To also enable mass calibration on supported lipid bilayers, membranes containing 0.01 mol % Biotinyl Cap PE were used. As linker for the attachment of biotinylated standard proteins, 2.5 nM divalent streptavidin were incubated for 10 min (RT) prior to the addition of 100 nM biotin labeled bovine albumin (#A8549, Sigma Aldrich, St. Louis, USA), biotinylated Pierce™ protein A (#29989, Thermo Fisher Scientific, Waltham, USA) or custom biotinylated aldolase (#28403842, Cytiva, Marlborough, USA). These concentrations correspond to 3-6 particles (median value) in the field of view for each standard protein, equivalent to a particle density of 0.09 to 0.19 µm^−2^.

For diffusion coefficient verification, 1.25 nM unconjugated tetravalent Streptavidin (#SNN1001, Thermo Fisher Scientific, Waltham, USA) were incubated for 10 min (RT) on a supported lipid membrane containing biotinylated lipids.

### Mass Photometry Landing Assay

To determine the nucleotide-dependent solution-state of MinD (175 nM) and MinDE (175 nM), all proteins were diluted in filtered (sterile syringe filters 0.45 *μ*m cellulose acetate membrane, VWR International, Radnor, USA) Min buffer in the presence of either 0.5 mM ADP or ATP. Measurements were performed in flow chambers and 50 *μ*L protein solution were flushed in five times consecutively, to collect sufficient landing events.

### *In Vitro* Reconstitution of Min Complexes on a Lipid Bilayer

#### MinD

For MSPT of single MinD complexes on the lipid interface, we added increasing protein concentrations (50, 75, 100, 125, 150, 175 and 200 nM, n = 3 flow chambers each) to a flow chamber with a supported lipid bilayer in the presence of 0.5 mM ATP in filtered Min buffer. The same experiments were performed likewise for MinD D40A. Videos of 0.5 mM ATP in Min buffer without protein were recorded as image background control (0.8 % of particles detected as compared to videos containing MinD).

#### MinDE

Co-reconstitution of MinDE was performed in a similar manner as for MinD, except for the addition of equimolar amounts of both reactants (50, 75, 100, 125, 150, 175 and 200 nM, n = 3 flow chambers each) to the sample chamber. For MinD D40AE, experiments were performed with 200 nM of each protein.

## Microscopy

Imaging of MinDE landing assays was performed on a custom-built interferometric scattering microscope described in ^17,54^ with a 445 nm laser diode for illumination and 635 nm for focus-stabilization. Image acquisition was controlled using custom-written software in Labview described in ^17^. Landing events were recorded at a frame rate of 1 kHz for 60 s per video. Videos were saved 5-fold frame-averaged (200 Hz effective frame rate) and 3-fold pixel-binned (70.2 nm effective pixel size).

SLB experiments were carried out on a commercial Refeyn One^MP^ mass photometer (Refeyn Ltd., Oxford, UK). Movies were acquired for either 45 s (landing assay of mass standards, Fig. 1) or 350 s (SLB assays) with the Aquire^MP^ (Refeyn Ltd., v2.3) software at a frame rate of 1 kHz. Movies were saved 5-fold frame-averaged (200 Hz effective frame rate) and 4-fold pixel-binned (84.4 nm effective pixel size).

## Data analysis

### Image Processing of Conventional MP Landing Assays

Videos of the proteins MinD, MinE and their equimolar mixture MinDE landing on glass cover slides were analysed with the software Discover^MP^ (version 2.1.0, Refeyn Ltd.), using a rolling background removal strategy^17^.

MinD monomers have a molecular mass of 33 kDa, which approaches the lower detection limit of the mass photometer. We have therefore systematically determined the optimal frame averaging factor *navg* and filter thresholds *T1* and *T2* for the detection of MinD monomers and dimers. To this end, we generated a semi-synthetic movie that used frames from a video recorded in a chamber with Min buffer alone to reconstruct the experimental background and added simulated PSFs (same as fitting model PSF) as landing events that had the expected scattering contrast of MinD monomers (contrast = 1.9 × 10^−3^) or MinD dimers (contrast = 3.5 × 10^−3^). We then varied *navg* as well as *T1* and *T2*, ran the analysis procedure, and evaluated the number of true positive and false positive detections. To determine the maximum number of true positive detections possible at the respective signal-to-noise ratio, we simulated 1000 frames with 100 landing events that were not allowed to spatially overlap closer than 12 pixels and 26 frames temporally. Based on the simulation, we chose *navg* = 12, *T1* = 1.1 and *T2* = 0.2 to process the experimental videos. Using these parameters, the number of true positive detections of monomers was 68.8 ± 6.6% (mean ± STD, n = 5 simulated videos), while the total of false positive detections was 9.4 ± 2.1%. For dimers, a simulation with the same parameters gave 95.8 ± 0.8% true positive detections and 4.2 ± 1.1% false positive ones.

### Mass Calibration for Landing Assays

Landing events imaged in three independent experiments per standard protein were pooled and kernel density estimations were calculated. For each standard protein, the contrast associated with a probability density maximum was determined and plotted against its nominal mass (see Supplementary Table 2). In case of alcohol dehydrogenase and β-amylase, the peak with higher contrast was used.

Error bars in Fig 1c represent the standard error of the peak position calculated from 10,000 bootstrap resamples.

#### Image Processing and Analysis Procedure for Single-Particle Tracking and Mass Determination of Biomolecules on SLBs

To detect and analyse diffusing biomolecules on a lipid bilayer, we applied a new image processing strategy that removed the dominant static scattering background, while conserving the shape of mobile features and displaying them on top of a shot-noise limited background. Around each frame, a pixel-wise temporal median image was calculated that only contains the features which did not move in the median period. This static background, calculated for each frame, was then removed from the respective frame with index i according to equation (1) and with n denoting the median half-size.

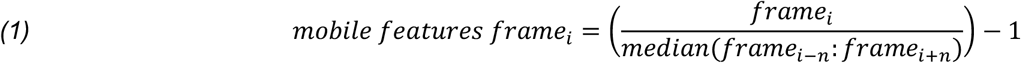

Through this process, moving objects in the mobile features frame appeared as undistorted PSFs and displayed iSCAT contrasts similar to those obtained for molecules in the conventional landing assay. We kindly received a custom-written Python script from Philipp Kukura and Gavin Young which automatised this image analysis procedure and we modified it to load and process videos in the Refeyn format (.mp files). To track individual proteins diffusing on supported lipid bilayers in the background-corrected movies, the Python script also included a single-particle tracking routine. For particle detection, a Laplace filter was applied to each frame (*scipy*.*ndimage*.*filters*.*gaussian_laplace* function), to suppress shot noise and highlight potential particles. The Laplace-filtered images were thresholded, such that only objects with at least the size of 53 kDa remained, a threshold that would preserve streptavidin as our smallest standard protein and MinD dimers. Local maxima were then found (*scipy*.*ndimage*.*filters*.*maximum_filter* function) as candidate pixels that contained a particle. Around each of these candidate pixels in the non-Laplace-filtered image, a region of interest (ROI) of 13 by 13 pixels (84.4 nm/pixel) was excised and fit by the model PSF used in the Discover^MP^ software to extract particle contrast and location at sub-pixel resolution. Particle locations were linked into trajectories using the *Trackpy* package (version 0.4.2, *link_df* function) ^55^. Only trajectories of particles with a lifetime of at least 5 frames (25 ms) were considered for subsequent analyses. Supplementary Table 1 summarises all parameters used for particle detection, fitting and trajectory linking.

#### Determination of Diffusion Coefficients

To determine translational diffusion coefficients from individual trajectories, a jump-distance analysis was performed as described in ^36^. Cumulative frequency distributions of jump distances for time lags from Δ1 to Δ4 frames were fitted globally (*scipy*.*optimize*.*least_squares*, Trust Region Reflective algorithm) with either one or two species depending on which model resulted in a reduced chi-squared statistic.

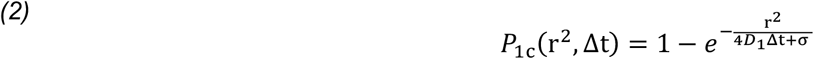

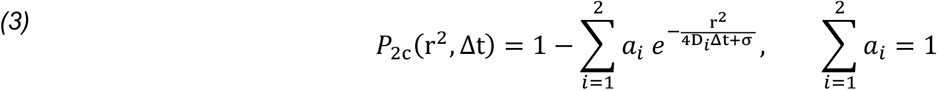

*D*_i_ and *a*_*i*_ are the diffusion coefficient and abundance of component *i, Δt* is the time lag and *σ* is an offset parameter taking into account localisation uncertainty and motion blur ^56^. In the case of a two-component model (~10% of all trajectories), an effective diffusion coefficient was calculated according to

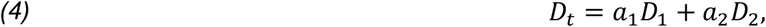

For comparison with the jump distance approach in the diffusion simulation (Supplementary Figure 2), diffusion coefficients were also estimated based on the mean squared displacement (MSD). Here, the first three or four non-trivial time-averaged MSD points were included in the linear regression depending on the trajectory length ^56^.

In both fit routines, the diffusion coefficient was restricted to values greater than 0.0001 µm^2^/s while the offset parameter σ was unconstrained. Trajectories where the diffusion coefficient returned by the fit hit the lower boundary were excluded from the analysis.

#### Kernel Density Estimation in 1D

For univariate distributions, kernel density estimations (KDEs) were computed with the Python package *KDEpy* (version 1.0.10, *FFTKDE* function) [https://github.com/tommyod/KDEpy], using a Gaussian kernel and the Improved Sheather-Jones plug-in bandwidth selector^57^.

#### Kernel Density Estimation in 2D

2D maps of the diffusion coefficient and mass were computed with the Python package *fastkde* (version 1.0.14, *fastkde*.*pdf* function) ^58^. The probability density for the diffusion coefficient was calculated in base 10 logarithmic space. Marginal distributions displayed on top and on the right of the graph were calculated by summing up the bivariate probability densities along the y- and x-axis, respectively.

The resulting 2D kernel densities were plotted as filled contours with either six (if an overlay of two conditions is shown) or eight linearly spaced levels. The opacity of the levels was reduced linearly from 100% for the highest to 0% for the lowest level, rendering the lowest level fully transparent.

#### Mass Calibration for MSPT

Analogous to the mass calibration for landing assays, peak contrasts were related to the nominal mass of streptavidin (Strep) or to the mass of streptavidin and the standard protein (Strep-ALD, Strep-BSA, Strep-prA). For the streptavidin-protein complexes, the peak with lowest contrast was excluded from analysis as it resembles streptavidin without an attached standard protein (blue crosses, Supplementary Fig. 4). The two other peak(s) with higher contrast were assigned to dimeric and tetrameric aldolase, monomeric and dimeric BSA and monomeric protein A (Supplementary Table 3).

Error bars in Fig 1c represent the standard error of the peak position calculated from 10,000 bootstrap resamples.

#### Simulated Videos of Diffusing Particles

To investigate the influence of extreme diffusion coefficients and high particle densities on the fidelity of our MSPT analysis, we implemented a Python routine that adds artificial diffusing particles to an experimental video of an empty SLB. Thereby, it was possible to control the number of particles in the field of view (FOV) and to individually set both the iSCAT contrast and the particles’ diffusion coefficient. In a first step, the trajectories of all particles were generated by choosing a random starting location within an area that is 4 times the video FOV size (x- and y-axes extended 2-fold). The travelled per-frame distance *Δs* in the × and y directions was drawn randomly from normal distributions with standard deviation *2DΔt* (equations 5).

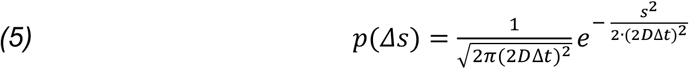

*D* is the diffusion coefficient converted into pixel dimensions and *Δt* the frame time. In order to keep the number of particles in the simulated area constant, we added periodic boundaries. Whenever a particle would cross a boundary of the simulated area, its location was shifted to the corresponding location at the opposite boundary. Trajectory coordinates that entered the FOV of the experimental video, i.e., 128 × 35 pixels, were populated with PSFs of a defined ratiometric contrast according to the Refeyn fitting model. Finally, the background-free ratiometric video of simulated PSFs was merged with a raw experimental iSCAT video of an empty SLB via pixel-wise calculation of

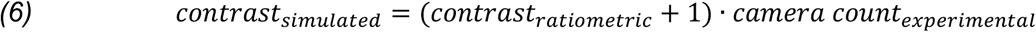

The resulting simulated videos were then analysed by our MSPT analysis routine in the same way as an experimental video with real particles.

#### Analysis of MinD and MinD40A Oligomerisation

Each trajectory was assigned to an apparent membrane protein density determined as the median of all trajectories detected during the trajectory’s lifetime, divided by the area of the FOV (31.9 µm^2^). For trajectory densities ranging from 0.03 to 0.8 µm^−2^ (corresponding to 1 to 27 particles in the FOV), the KDE of each mass distribution was calculated. The densities were binned in an overlapping manner, such that all trajectories were selected that had trajectory numbers within a range of plus or minus two (e.g. at a density of 0.1 µm^−2^, corresponding to 4 trajectories/FOV, trajectories with membrane coverage between 2 and 6 trajectories/FOV were pooled). Each KDE was fit with a linear combination of monomer to hexamer mass distributions that were inferred from single component simulations of MinD or MinD D40A at varying particle densities (Supplementary Fig. 9). The mass distributions for each oligomer and density were selected based on the number of localised particles in the experiment as a certain fraction of particles (~40%) is lost in the trajectory linking and filtering (minimum length of 5 frames) process (Supplementary Fig. 12). Additionally, the sum of contributions of the six components was constrained to add up to unity in the least-squares sense. Prior to fitting, the simulated mass distributions were smoothed using a moving average with a window length of 4.5 kDa. To estimate the uncertainty of the extracted abundances of the components, the dataset at each trajectory density was split randomly into three samples before fitting each subset individually. This procedure was repeated 10 times and the average standard deviation of the persplit results was calculated. Unless otherwise stated, the reported particle densities throughout the manuscript correspond to trajectory densities (i.e. linked particle densities) as a more robust means to estimate the current membrane coverage (in contrast to localised particle densities; see Supplementary Figs. 7, 12).

#### Step Detection

To detect mass change points during trajectories, we employed a MATLAB (R2020a, The MathWorks, Natick, MA) implementation of the Kalafut-Visscher algorithm^59^ described in^47^. While the algorithm has no parameters except the time series itself, we noticed that the results depended on the length of the time series and the position of potential change points if they were located close to the beginning or the end of a trajectory. Thus, change points were more likely to be inserted, the shorter a time series was and the closer a point was to the boundaries of the time series. To address this ambiguity, we concatenated the trajectories of a specific imaging condition (e.g. of all videos containing MinD, Supplementary Fig. 14a) and divided this concatenated series into *n* subsets of equal length *l*, which were then analysed with the step detection algorithm (Supplementary Fig. 14b). This procedure enabled an equal treatment of trajectories of different lengths and avoided bias in change point detection at the beginning and the end of trajectories. Additionally, step identification was repeated *l – 1* times shifting the start point of the linked time series circularly by one increment in each iteration. This second procedure ensured that the detected steps would not depend on the start points of a subset. From the *l* outputs generated in total, the relative significance of a step was reflected by the fraction *f* of iterations, in which a change point was found at a particular location, ranging from 0 (never) to 1 (in every iteration, Supplementary Fig. 14c).

The choice of fraction *f* as well as the length *l* is somewhat arbitrary yet interconnected. Increasing *l* leads to fewer detected steps which can be compensated for by reducing *f*. Since the longest trajectory in all datasets has a length of 558 frames and an increasing length leads to a finer grating of the tunable parameter *f*, we set *l* to 1000 frames. By visual inspection, we chose a fraction *f* of 0.25 as an appropriate tradeoff between under- and over-identification of steps considering the noise level in our trajectories (Supplementary Fig. 14d).

#### Subunit Attachment/Detachment During Membrane Diffusion and Release of Complexes

The concatenated mass traces and step-fitting results were split into their original trajectories (Supplementary Fig. 14e). For each trajectory, step heights (equivalent to mass changes) and dwell lengths (equivalent to time intervals without a change of mass) were extracted and categorised into three classes (Fig. 3b). Depending on whether the sign of the succeeding mass change is positive or negative, the preceding level is classified as an attachment (at) or as a detachment plateau (dt), respectively. The last plateau in a trajectory prior to the dissociation of the whole particle from the membrane is regarded separately as release (rl). If no step is detected in the mass time series as it was the case in ~90% of all trajectories, the single plateau is also classified as release (rl). Particles leaving the FOV have been excluded from the analysis. Therefore, it can be assumed that the end of a trajectory represents the particle’s release from the membrane. Generally, the mass which is initially binding to the membrane is not included in the list of steps for attachment or detachment events.

The distribution of mass changes during the trajectory as well as the distribution of complex mass released from the membrane for different densities are displayed as univariate kernel density estimates. To characterise the residence times of different oligomers, histograms of the observed dwell lengths were generated for plateaus with masses that correspond to dimeric (66 ± 17 kDa) or tetrameric (132 ± 17 kDa) MinD complexes. For each variant the average dwell times were calculated according to equation *(7)*

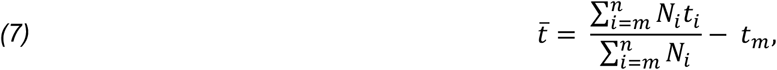

where *N*_i_ is the total number of plateaus with dwell time *t*_i_, and *t*_m_ represents the most frequent dwell time. Due to the noise level of mass detections in our trajectories, short dwell times could not be quantitatively detected. Hence, dwell times shorter than 5 frames (25 ms) were excluded from the analysis.

The standard error of the mean dwell time was estimated from 10,000 bootstrap resamples.

## Acknowledgments

We gratefully thank the group of Philipp Kukura at the University of Oxford for hosting early experiments and Gavin Young for sharing image processing and particle tracking code. We thank Max Felix Hantke, Josiah Kane, and the rest of the Refeyn software team for providing mass photometry image analysis code and their support. We thank Daniel Bollschweiler (Cryo-EM MPIB Core Facility) for the initial introduction to the commercial Refeyn One^MP^ mass photometer, the MPIB Biochemistry Core Facility (Recombinant Protein Production) and Kerstin Andersson for assistance with protein purification, S. Bauer for assistance with vesicle preparation and Florian Stehr for helpful discussions. T.H. and P.S. acknowledge funding through the Deutsche Forschungsgemeinschaft (DFG, German Research Foundation) - Project-ID 201269156 – SFB 1032 (A09) and B.R. and P.S. through Project 269423233 – TRR 174. B.R. was supported by a DFG fellowship through the Graduate School of Quantitative Biosciences Munich (QBM). N.H. was supported by a DFG return grant HU 2462/3-1. P.S. acknowledges support through the research network MaxSynBio via the joint funding initiative of the German Federal Ministry of Education and Research (BMBF) and the Max Planck Society.

## Author Contributions

T.H., F.S., N.H. contributed equally to this work. T.H., N.H., F.S. and P.S. conceived the study. T.H. designed and performed all experiments. B.R. designed and purified proteins. F.S. and N.H. analysed the data. T.H., F.S. and N.H. wrote the manuscript draft. All authors discussed and interpreted results. All authors revised, reviewed and approved the manuscript.

## Data Availability Statement

The datasets generated and analysed during the current study are available from the corresponding authors on reasonable request.

## Code Availability Statement

The custom-written Python script for particle detection and particle simulation (both as binary executable) as well as particle analysis (source code accessible) is available from the corresponding authors on reasonable request.

## Abbreviations

(ATP): adenosine-5’-triphosphate
(ADP): adenosine-5’-diphosphate
(BSA): bovine serum albumin
(18:1 Biotinyl Cap PE): 1,2-dioleoyl-sn-glycero-3-phosphoethanolamine-N-cap biotinyl
(DOPC): dioleoyl-sn-glycero-3-phosphocholine
(DOPG): dioleoyl-sn-glycero-3-phosphoglycerol
(HRP): horseradish peroxidase
(iSCAT): interferometric scattering
(P-loop): phosphate binding loop
(PSF): point spread function
(SPT): single-particle tracking
(SUV): small unilamellar vesicle
(SLB): supported lipid bilayer
(MP): mass photometry
(MSPT): mass-sensitive particle tracking
(MTS): membrane targeting sequence

## References

1. Gonzalez, L. & Scheller, R. H. Regulation of membrane trafficking: Structural insights from a Rab/effector complex. Cell vol. 96 755–758 (1999).

2. Cho, W. & Stahelin, R. V. Membrane-protein interactions in cell signaling and membrane trafficking. Annual Review of Biophysics and Biomolecular Structure vol. 34 119–151 (2005).

3. Bagheri, Y., Ali, A. A. & You, M. Current Methods for Detecting Cell Membrane Transient Interactions. Frontiers in Chemistry vol. 8 (2020).

4. Miller, H., Zhou, Z., Shepherd, J., Wollman, A. J. M. & Leake, M. C. Single-molecule techniques in biophysics: A review of the progress in methods and applications. Reports Prog. Phys. 81, 024601 (2018).

5. Manzo, C. & Garcia-Parajo, M. F. A review of progress in single particle tracking: From methods to biophysical insights. Reports on Progress in Physics vol. 78 (2015).

6. Gelles, J., Schnapp, B. J. & Sheetz, M. P. Tracking kinesin-driven movements with nanometre-scale precision. Nature 331, 450–453 (1988).

7. Funatsu, T., Harada, Y., Tokunaga, M., Saito, K. & Yanagida, T. Imaging of single fluorescent molecules and individual ATP turnovers by single myosin molecules in aqueous solution. Nature 374, 555–559 (1995).

8. Schmidt, T., Schütz, G. J., Baumgartner, W., Gruber, H. J. & Schindler, H. Imaging of single molecule diffusion. Proc. Natl. Acad. Sci. U. S. A. 93, 2926–2929 (1996).

9. Taylor, R. W. et al.. Interferometric scattering microscopy reveals microsecond nanoscopic protein motion on a live cell membrane. Nat. Photonics 13, 480–487 (2019).

10. Kukura, P. et al.. High-speed nanoscopic tracking of the position and orientation of a single virus. Nat. Methods 6, 923–927 (2009).

11. Jacobsen, V., Stoller, P., Brunner, C., Vogel, V. & Sandoghdar, V. Interferometric optical detection and tracking of very small gold nanoparticles at a water-glass interface. Opt. Express 14, 405 (2006).

12. Ueno, H. et al.. Simple dark-field microscopy with nanometer spatial precision and microsecond temporal resolution. Biophys. J. 98, (2010).

13. Loose, M., Fischer-Friedrich, E., Herold, C., Kruse, K. & Schwille, P. Min protein patterns emerge from rapid rebinding and membrane interaction of MinE. Nat. Struct. Mol. Biol. 18, 577–583 (2011).

14. Ha, T. & Tinnefeld, P. Photophysics of fluorescent probes for single-molecule biophysics and super-resolution imaging. Annual Review of Physical Chemistry vol. 63 595–617 (2012).

15. Garcia-Parajo, M. F., Segers-Nolten, G. M. J., Veerman, J. A., Greve, J. & Van Hulst, N. F. Real-time light-driven dynamics of the fluorescence emission in single green fluorescent protein molecules. Proc. Natl. Acad. Sci. U. S. A. 97, 7237–7242 (2000).

16. Lindfors, K., Kalkbrenner, T., Stoller, P. & Sandoghdar, V. Detection and spectroscopy of gold nanoparticles using supercontinuum white light confocal microscopy. Phys. Rev. Lett. 93, (2004).

17. Young, G. et al.. Quantitative mass imaging of single biological macromolecules. Science 360, 423–427 (2018).

18. Piliarik, M. & Sandoghdar, V. Direct optical sensing of single unlabelled proteins and super-resolution imaging of their binding sites. Nat. Commun. 5, (2014).

19. de Boer, P. A. J., Crossley, R. E. & Rothfield, L. I. A division inhibitor and a topological specificity factor coded for by the minicell locus determine proper placement of the division septum in E. coli. Cell 56, 641–649 (1989).

20. Raskin, D. M. & de Boer, P. A. MinDE-dependent pole-to-pole oscillation of division inhibitor MinC in Escherichia coli. J. Bacteriol. 181, 6419–24 (1999).

21. Raskin, D. M. & De Boer, P. A. J. Rapid pole-to-pole oscillation of a protein required for directing division to the middle of Escherichia coli. Proc. Natl. Acad. Sci. U. S. A. 96, 4971–4976 (1999).

22. Szeto, T. H., Rowland, S. L., Rothfield, L. I. & King, G. F. Membrane localization of MinD is mediated by a C-terminal motif that is conserved across eubacteria, archaea, and chloroplasts. Proc. Natl. Acad. Sci. U. S. A. 99, 15693–15698 (2002).

23. Hu, Z., Gogol, E. P. & Lutkenhaus, J. Dynamic assembly of MinD on phospholipid vesicles regulated by ATP and MinE. Proc. Natl. Acad. Sci. 99, 6761–6766 (2002).

24. Hsieh, C. W. et al.. Direct MinE-membrane interaction contributes to the proper localization of MinDE in E. coli. Mol. Microbiol. 75, 499–512 (2010).

25. Shih, Y. L. et al.. The N-terminal Amphipathic Helix of the topological specificity factor mine is associated with shaping membrane curvature. PLoS One 6, e21425 (2011).

26. Loose, M., Fischer-Friedrich, E., Ries, J., Kruse, K. & Schwille, P. Spatial regulators for bacterial cell division self-organize into surface waves in vitro. Science 320, 789– 792 (2008).

27. Miyagi, A., Ramm, B., Schwille, P. & Scheuring, S. High-speed atomic force microscopy reveals the inner workings of the MinDE protein oscillator. Nano Lett. 18, 288–296 (2018).

28. Ayed, S. H. et al.. Dissecting the role of conformational change and membrane binding by the bacterial cell division regulator MinE in the stimulation of MinD ATPase activity. J. Biol. Chem. 292, 20732–20743 (2017).

29. Ghosal, D., Trambaiolo, D., Amos, L. A. & Löwe, J. MinCD cell division proteins form alternating co-polymeric cytomotive filaments. Nat. Commun. 5, 5341 (2014).

30. Ye, W. et al.. Plasmonic Nanosensors Reveal a Height Dependence of MinDE Protein Oscillations on Membrane Features. J. Am. Chem. Soc. 140, (2018).

31. Heermann, T., Ramm, B., Glaser, S. & Schwille, P. Local Self-Enhancement of MinD Membrane Binding in Min Protein Pattern Formation. J. Mol. Biol. 432, 3191–3204 (2020).

32. Kretschmer, S., Zieske, K. & Schwille, P. Large-scale modulation of reconstituted Min protein patterns and gradients by defined mutations in MinE’s membrane targeting sequence. PLoS One 12, 1–16 (2017).

33. Ortega Arroyo, J. et al. Label-free, all-optical detection, imaging, and tracking of a single protein. Nano Lett. 14, 2065–2070 (2014).

34. Mosby, L. S. et al.. Myosin II Filament Dynamics in Actin Networks Revealed with Interferometric Scattering Microscopy. Biophys. J. 118, 1946–1957 (2020).

35. Howarth, M. et al.. A monovalent streptavidin with a single femtomolar biotin binding site. Nat. Methods 3, 267–273 (2006).

36. Weimann, L. et al.. A Quantitative Comparison of Single-Dye Tracking Analysis Tools Using Monte Carlo Simulations. PLoS One 8, (2013).

37. Johansson, B., Höök, F., Klenerman, D. & Jönsson, P. Label-free measurements of the diffusivity of molecules in lipid membranes. ChemPhysChem 15, 486–491 (2014).

38. Horton, M. R., Höfling, F., Rädler, J. O. & Franosch, T. Development of anomalous diffusion among crowding proteins. Soft Matter 6, 2648–2656 (2010).

39. Horton, M. R., Reich, C., Gast, A. P., Rädler, J. O. & Nickel, B. Structure and dynamics of crystalline protein layers bound to supported lipid bilayers. Langmuir 23, 6263–6269 (2007).

40. Stellmacher, L. et al.. Acid-Base Catalyst Discriminates between a Fructose 6-Phosphate Aldolase and a Transaldolase. ChemCatChem 7, 3140–3151 (2015).

41. Hu, Z. & Lutkenhaus, J. Topological regulation of cell division in E. coli: Spatiotemporal oscillation of MinD requires stimulation of its ATPase by MinE and phospholipid. Mol. Cell 7, 1337–1343 (2001).

42. Ramm, B. et al.. The MinDE system is a generic spatial cue for membrane protein distribution in vitro. Nat. Commun. 9, (2018).

43. Wu, W., Park, K. T., Holyoak, T. & Lutkenhaus, J. Determination of the structure of the MinD-ATP complex reveals the orientation of MinD on the membrane and the relative location of the binding sites for MinE and MinC. Mol. Microbiol. 79, 1515–1528 (2011).

44. Park, K. T., Wu, W., Lovell, S. & Lutkenhaus, J. Mechanism of the asymmetric activation of the MinD ATPase by MinE. Mol. Microbiol. 85, 271–281 (2012).

45. Evans, E. & Sackmann, E. Translational and rotational drag coefficients for a disk moving in a liquid membrane associated with a rigid substrate. J. Fluid Mech. 194, 553–561 (1988).

46. Khmelinskaia, A., Mücksch, J., Petrov, E. P., Franquelim, H. G. & Schwille, P. Control of Membrane Binding and Diffusion of Cholesteryl-Modified DNA Origami Nanostructures by DNA Spacers. Langmuir 34, 14921–14931 (2018).

47. Little, M. A. et al.. Steps and bumps: Precision extraction of discrete states of molecular machines. Biophys. J. 101, 477–485 (2011).

48. Park, K. T. et al.. The min oscillator uses MinD-dependent conformational changes in MinE to spatially regulate cytokinesis. Cell 146, 396–407 (2011).

49. Vecchiarelli, A. G. et al.. Membrane-bound MinDE complex acts as a toggle switch that drives Min oscillation coupled to cytoplasmic depletion of MinD. Proc. Natl. Acad. Sci. U. S. A. 113, E1479–E1488 (2016).

50. Ghasriani, H. & Goto, N. K. Regulation of symmetric bacterial cell division by MinE What is the role of conformational dynamics? Commun. Integr. Biol. 4, 101–103 (2011).

51. Lackner, L. L., Raskin, D. M. & De Boer, P. A. J. ATP-dependent interactions between Escherichia coli Min proteins and the phospholipid membrane in vitro. J. Bacteriol. 185, 735–749 (2003).

52. Ramm, B., Glock, P. & Schwille, P. In vitro reconstitution of self-organizing protein patterns on supported lipid bilayers. J. Vis. Exp. 1–13 (2018) doi:10.3791/58139.

53. Howarth, M. & Ting, A. Monovalent streptavidin expression and purification. Protoc. Exch. (2008) doi:10.1038/nprot.2008.81.

54. Cole, D., Young, G., Weigel, A., Sebesta, A. & Kukura, P. Label-Free Single-Molecule Imaging with Numerical-Aperture-Shaped Interferometric Scattering Microscopy. ACS Photonics 4, 211–216 (2017).

55. Allan, D. et al.. soft-matter/trackpy: Trackpy v0.4.2. (2019) doi:10.5281/ZENODO.3492186.

56. Michalet, X. Mean square displacement analysis of single-particle trajectories with localization error: Brownian motion in an isotropic medium. Phys. Rev. E - Stat. Nonlinear, Soft Matter Phys. 82, 041914 (2010).

57. Botev, Z. I., Grotowski, J. F. & Kroese, D. P. Kernel density estimation via diffusion. Ann. Stat. 38, 2916–2957 (2010).

58. O’Brien, T. A., Kashinath, K., Cavanaugh, N. R., Collins, W. D. & O’Brien, J. P. A fast and objective multidimensional kernel density estimation method: FastKDE. Comput. Stat. Data Anal. 101, 148–160 (2016).

59. Kalafut, B. & Visscher, K. An objective, model-independent method for detection of non-uniform steps in noisy signals. Comput. Phys. Commun. 179, 716–723 (2008).

