## Supplementary Information for "Mass-sensitive particle tracking (MSPT) to elucidate the membrane-associated MinDE reaction cycle"

**Title**

### Supplementary Figures

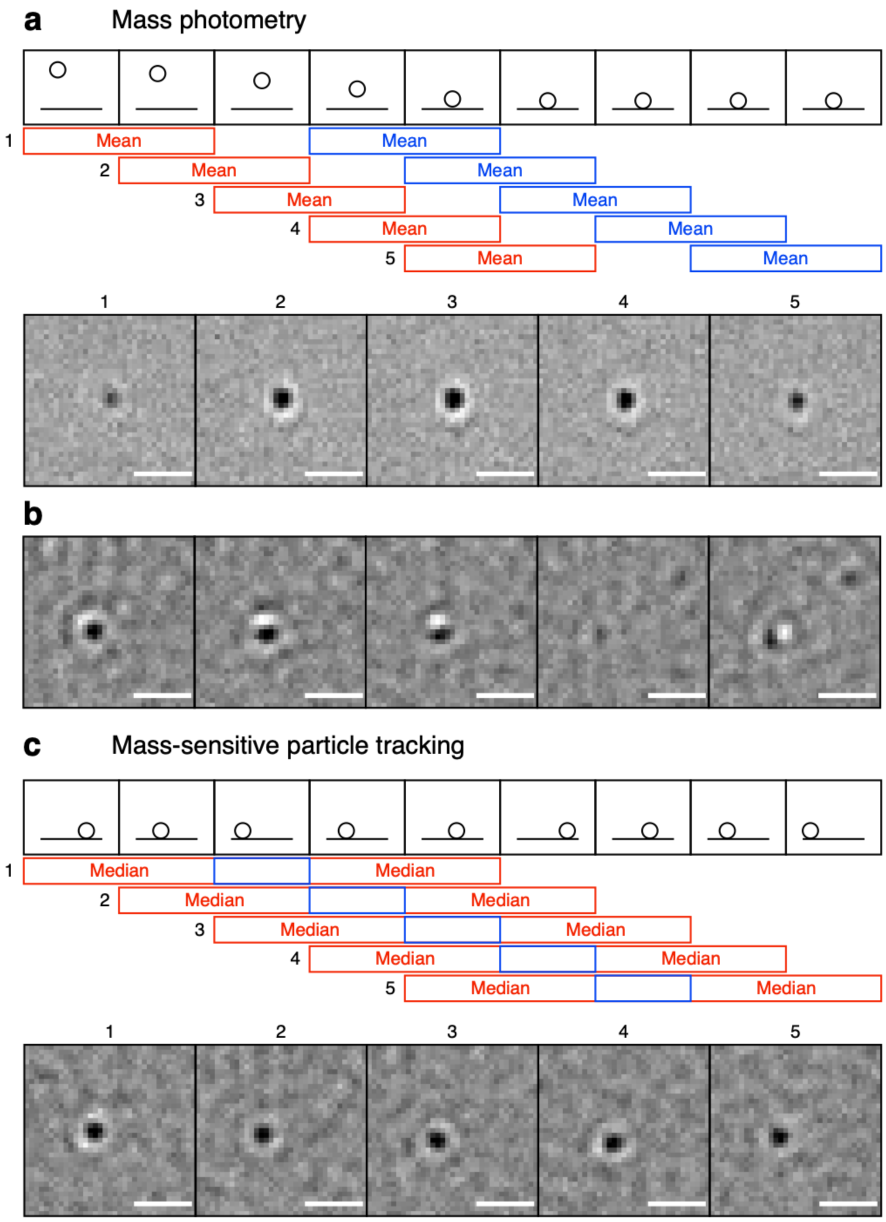


**Supplementary Figure 1 - Illustration and comparison of image processing procedures. (a)** In mass photometry (MP), landing molecules are visualised through gradually fading their PSF in and out of the video. Top: Schematic representation of a frame sequence of a landing particle. Middle: Mean of pixel intensities of blue frames is divided by the mean pixel intensities of red frames. Bottom: Ratiometric frames obtained for a landing event of β-amylase on glass applying this image-processing approach. **(b)** Ratiometric frames obtained for a biotin-aldolase-streptavidin complex diffusing on a biotinylated supported lipid bilayer using the processing scheme in (a). Note the distorted and occasionally disappearing PSF. **(c)** For mass-sensitive particle tracking (MSPT), mobile features are visualised by removing background that contains only static features. Top: Schematic representation of a frame sequence of a randomly moving particle on a surface. Middle: Pixel intensities of the blue central frame are divided by the median of pixel intensities of surrounding red frames. Bottom: Ratiometric frames obtained for the same biotin-aldolase-streptavidin complex as in (b) using the image-processing approach in (c). Scale bars: 1 µm.

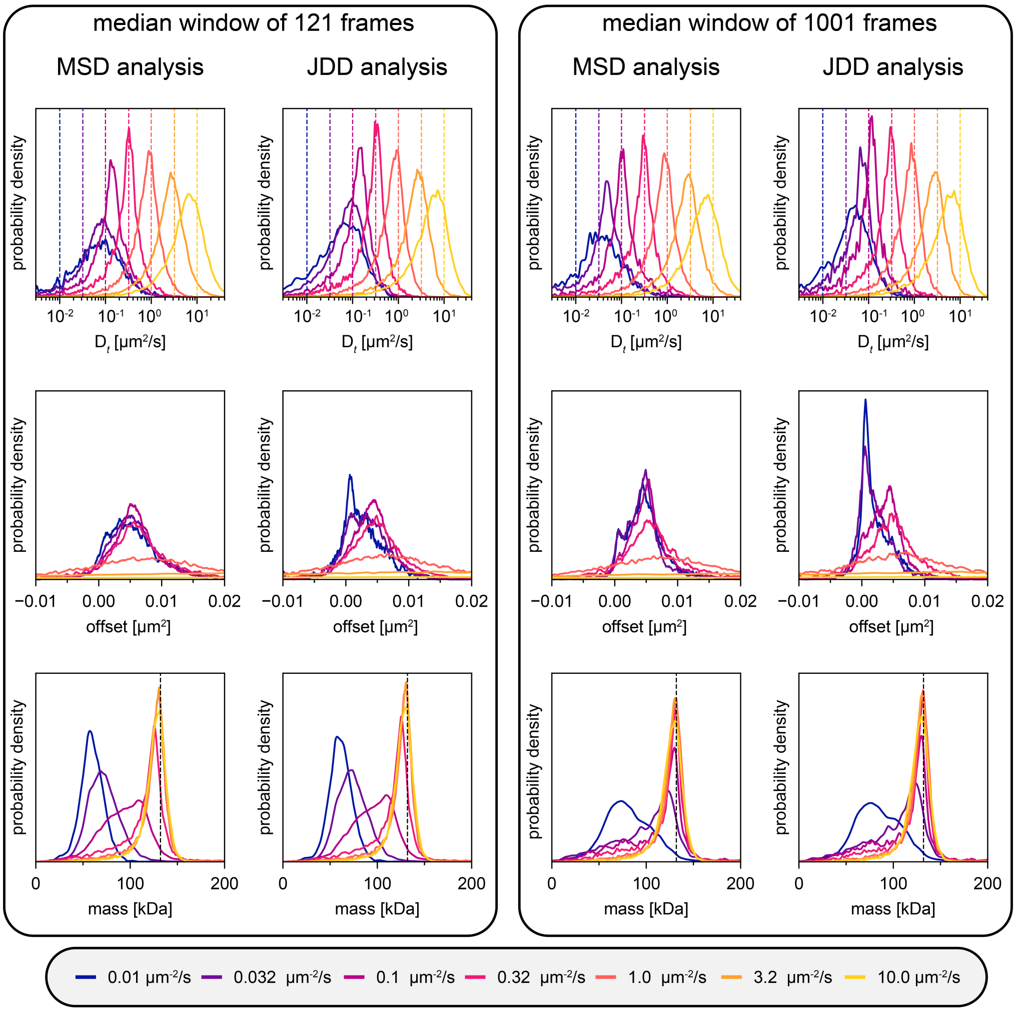


**Supplementary Figure 2 – Comparison of MSD and JDD performance using simulated MSPT videos with varying diffusion coefficients.** Both the left and right panel each display the distribution of determined diffusion coefficients (top row), the fitted offset corresponding to the particle localisation uncertainty (middle row) and the extracted mass distribution (bottom row) for MSD and JDD analysis. Simulated videos used for particle tracking were processed either with a median window of 121 frames (left panel) or with a median window of 1001 frames (right panel). In the top row, dashed lines indicate the input diffusion coefficient and the distributions in the corresponding colour show the diffusion coefficients extracted from the respective simulated videos. In the bottom row, the dashed line indicates the input mass (i.e. input particle contrast). The average particle density in the videos varied between 0.06 and 0.16 µm$^{-2}$. For the number of analysed trajectories, please refer to Supplementary Table 5.

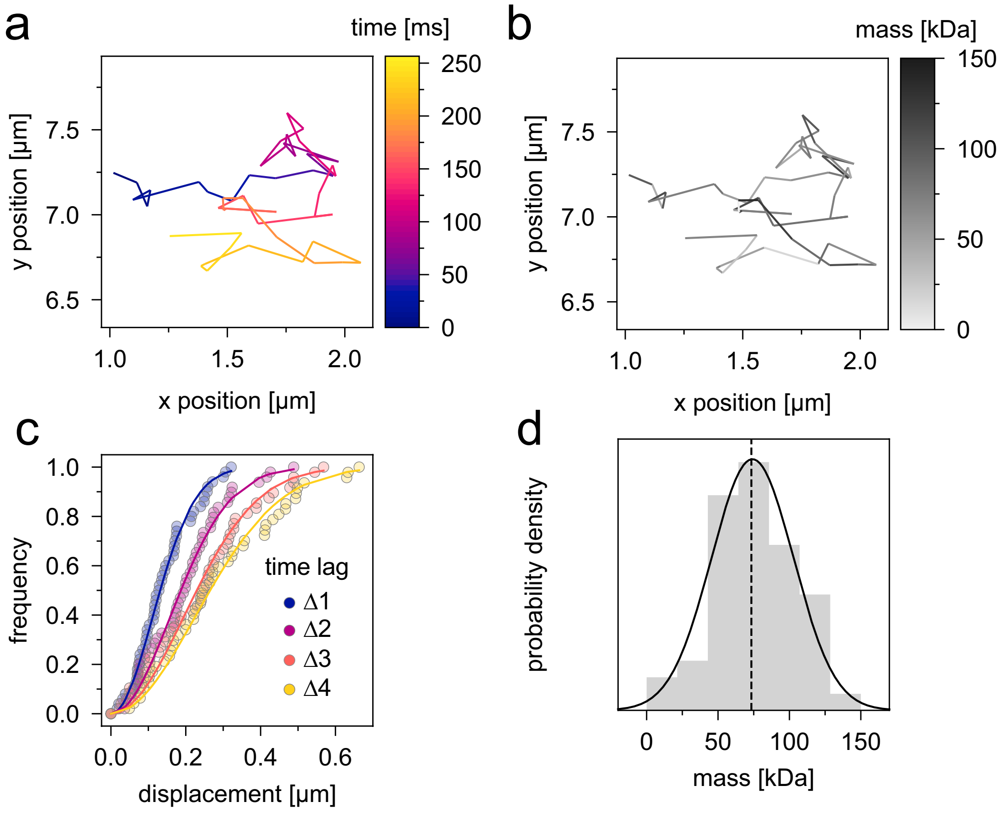


**Supplementary Figure 3 - Single-particle characterisation of the diffusion of membrane-attached streptavidin by MSPT. (a, b)** Representation of a reconstructed particle trajectory colour-coded by time (**a**) or mass (**b**). **(c)** Cumulative distribution of displacements for time lags from Δ1-4 frames globally fitted with an equation describing two-dimensional Brownian motion (see methods). **(d)** Histogram of the masses detected in each frame along the trajectory (grey) and its approximation with a normal distribution (black line). The dashed line represents the median.

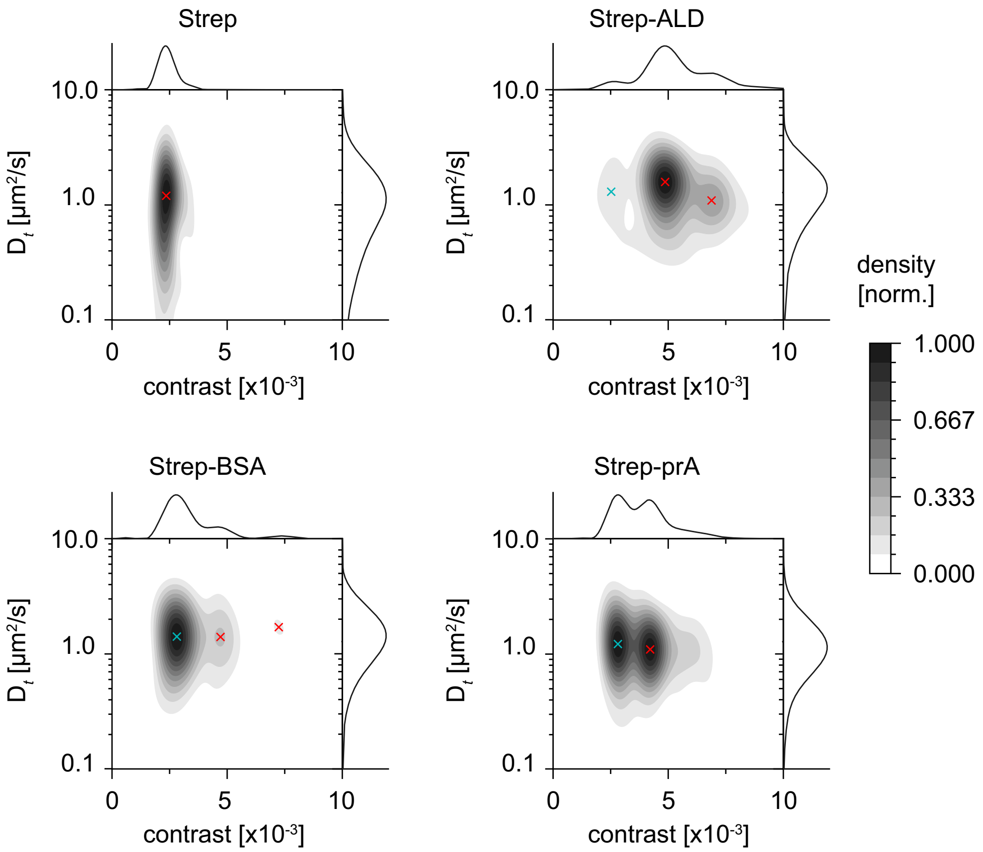


**Supplementary Figure 4 - 2D maps of diffusion coefficient and mass for biotinylated MSPT standard proteins attached via divalent streptavidin to biotinylated lipids on a supported lipid bilayer.** Only particles with a track length of at least 10 frames were included. Oligomer states included in the MSPT mass calibration are highlighted with a red x. A turquoise x mark peaks that were not considered for mass calibration because they represent unbound divalent streptavidin. Trajectory numbers: divalent streptavidin, Strep, n = 16,699; divalent streptavidin with biotinylated aldolase, Strep-ALD, n = 16,727; divalent streptavidin with biotinylated bovine serum albumin, Strep-BSA, n = 8,842 and divalent streptavidin with biotinylated protein A, Strep-prA, n = 22,424. Marginal probability distributions of both molecular mass and diffusion coefficient are presented at the top and right, respectively.

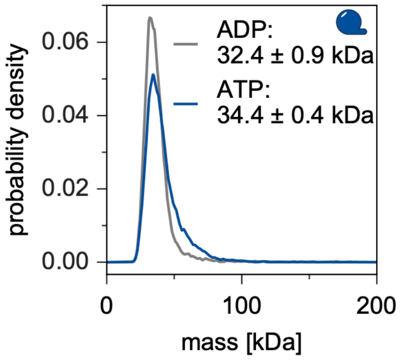


**Supplementary Figure 5 - The MinD monomer state is preserved in solution.** Mass distributions of 175 nM MinD in the presence of 0.5 mM ATP (blue line; n = 16,013 particles) or 0.5 mM ADP (grey line; n = 7,001 particles) determined by mass photometry.

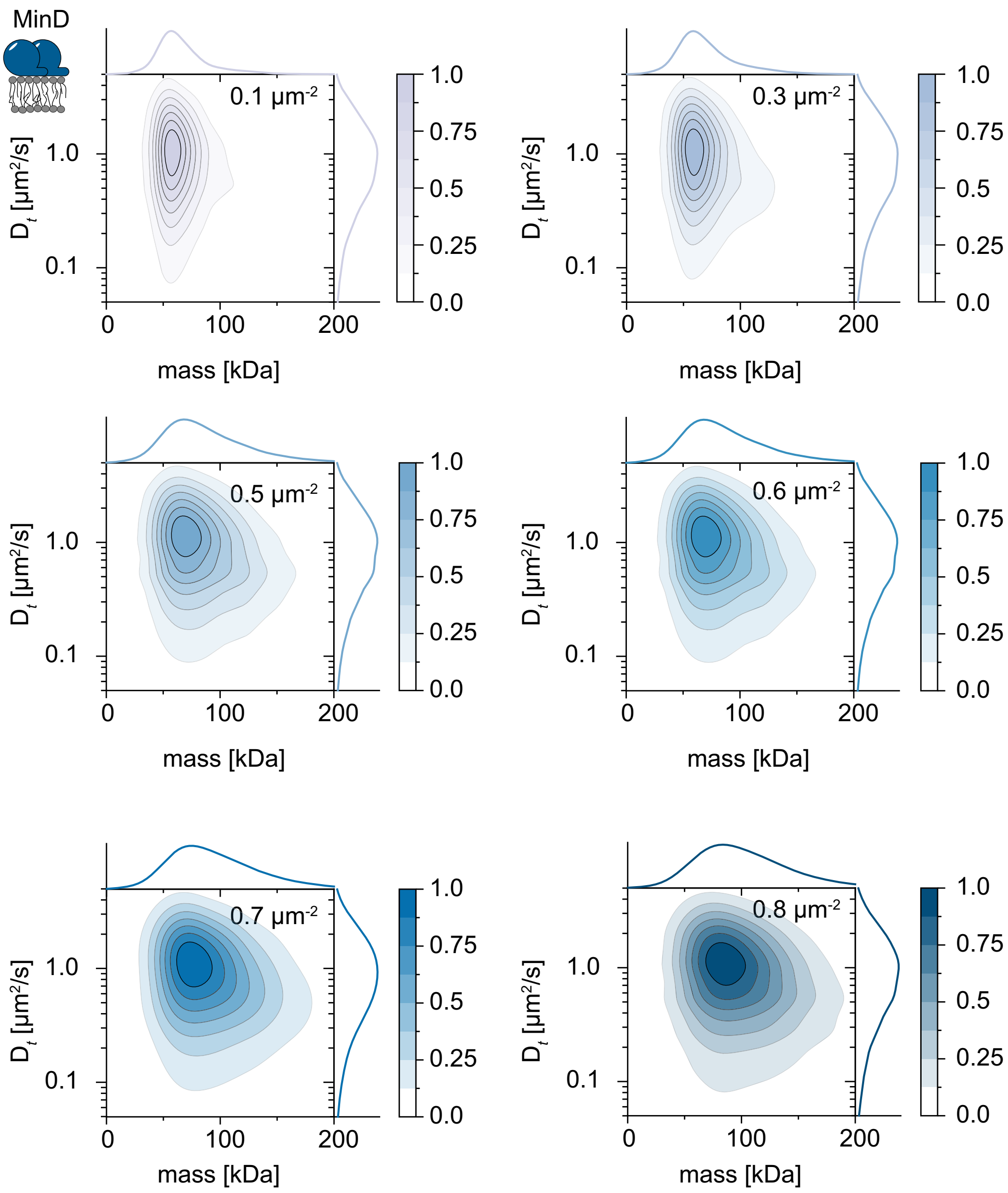


**Supplementary Figure 6 - 2D maps of diffusion coefficient and mass for membrane-bound MinD particles.** 2D kernel density estimation of diffusion coefficient versus mass for membrane-attached MinD at particle densities of 0.1 µm$^{-2}$ (light mauve, n = 117,086 trajectories), 0.3 µm$^{-2}$ (mauve, n = 169,957 trajectories), 0.5 µm$^{-2}$ (light blue, n = 256,404 trajectories), 0.6 µm$^{-2}$ (blue, n = 256,404 trajectories), 0.7 µm$^{-2}$ (dark blue, n = 282,414 trajectories) and 0.8 µm$^{-2}$ (midnight blue, n = 152,685 trajectories). Marginal probability distributions of both molecular mass and diffusion coefficient are presented at the top and right, respectively.

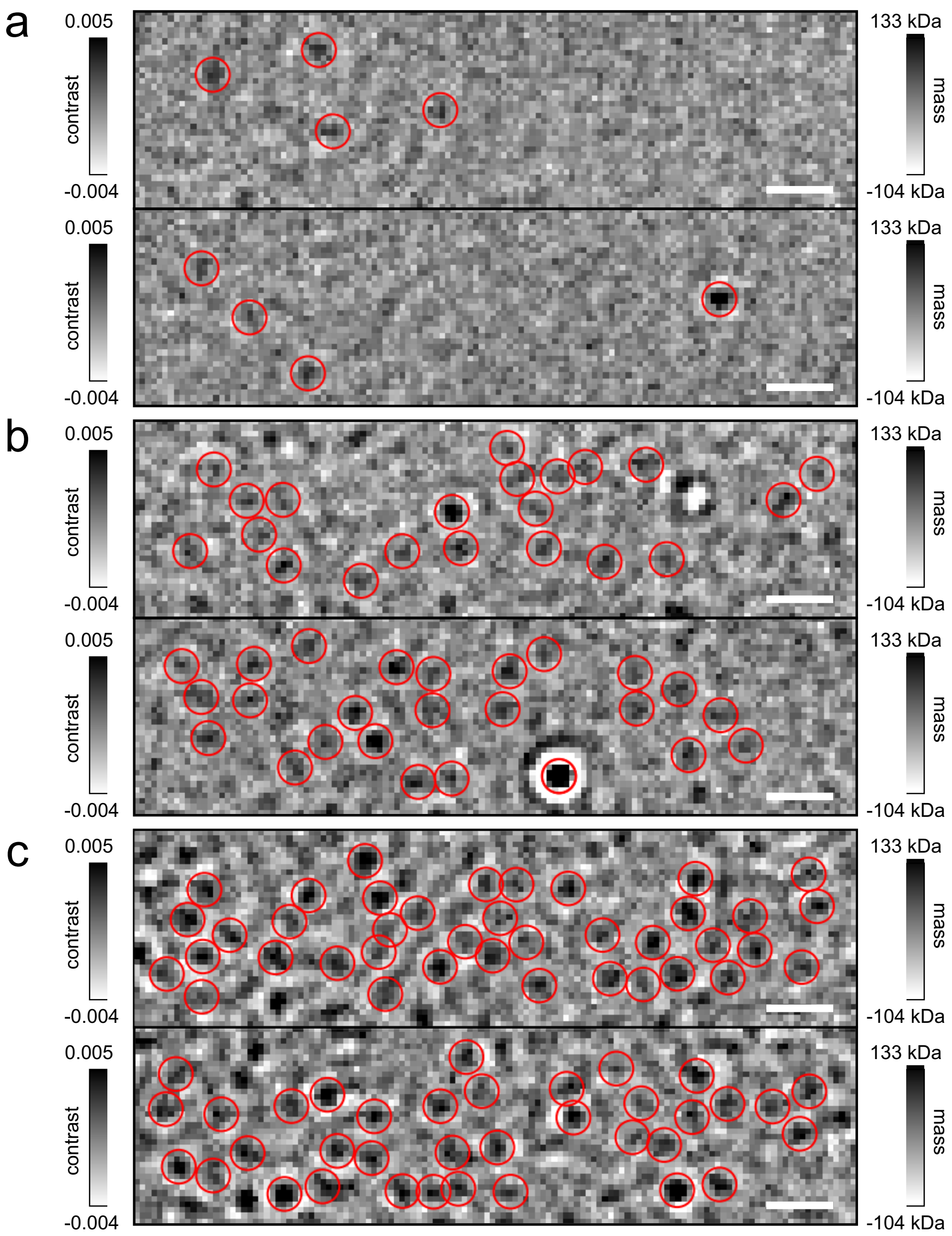


**Supplementary Figure 7 - Representative iSCAT images of diffusing MinD particles on a supported lipid bilayer at three different particle densities.** Detected and fitted MinD particles (i.e. localised particles) are highlighted through red circles at three different particle densities, 0.1 µm$^{-2}$ **(a)**, 0.5 µm$^{-2}$ **(b)** and 0.8 µm$^{-2}$ **(c)**. Note, that some candidates will be lost in the process of trajectory linking. Contrast range: black, 0.005 ≙ 133 kDa; white, -0.004 ≙-104 kDa. Scale bars: 1 µm.

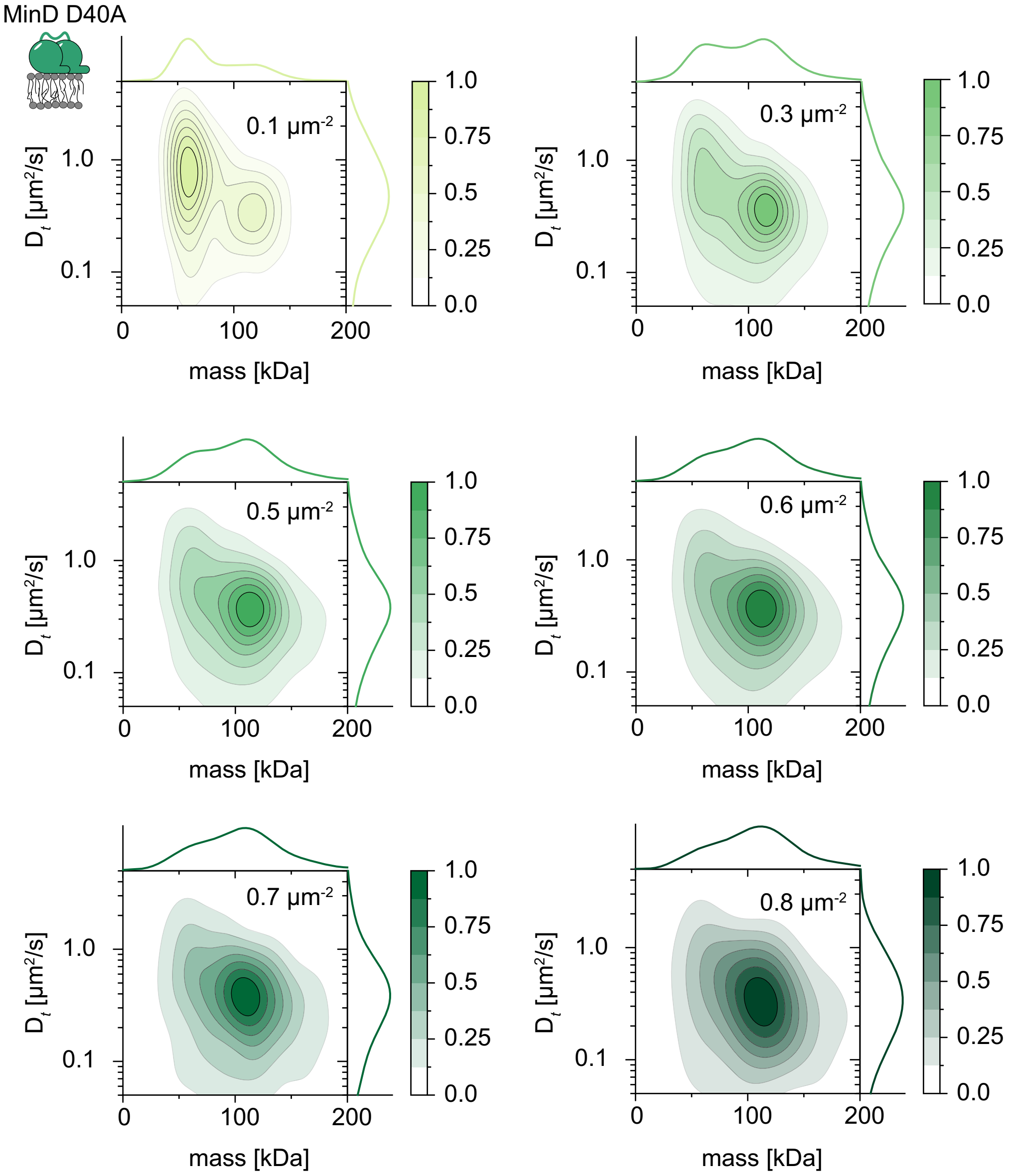


**Supplementary Figure 8 - 2D maps of diffusion coefficient and mass for membrane-bound MinD D40A particles.** 2D kernel density estimation of diffusion coefficient versus mass for membrane-attached MinD D40A at particle densities of 0.1 $\mu m^{-2}$ (lime, n = 7,831 trajectories), 0.3 $\mu m^{-2}$ (fern, n = 72,872 trajectories), 0.5 $\mu m^{-2}$ (emerald, n = 72,529 trajectories), 0.6 $\mu m^{-2}$ (sea green, n = 42,280 trajectories), 0.7 $\mu m^{-2}$ (seaweed, n = 12,172 trajectories) and 0.8 $\mu m^{-2}$ (viridian, n = 3,150 trajectories). Marginal probability distributions of both molecular mass and diffusion coefficient are presented at the top and right, respectively.

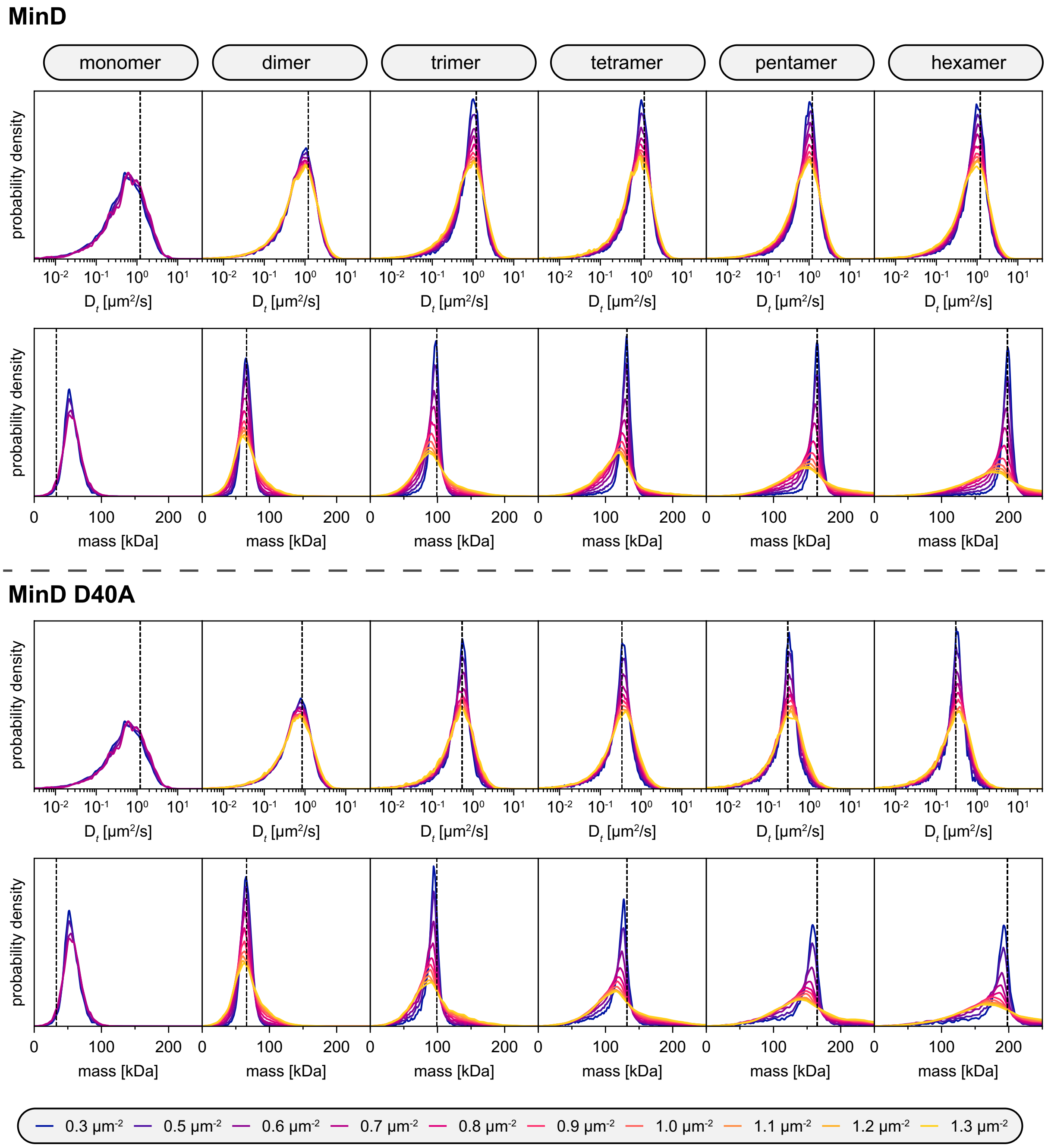


**Supplementary Figure 9 – Effect of high particle densities on extracted diffusion coefficient and particle mass in simulated MSPT videos.** Distributions of determined diffusion coefficients (top row) and extracted masses (bottom row) for videos with simulated particles corresponding to MinD and MinD D40A oligomers. The simulation was set up with particles that had a constant diffusion coefficient of 1.2 µm$^2$ s$^{-1}$ and iSCAT contrasts corresponding to a MinD monomer (33 kDa), a dimer (66 kDa), a trimer (99 kDa), a tetramer (132 kDa), a pentamer (165 kDa) and a hexamer (198 kDa). Simulations of MinD D40A particles were set up with identical masses as for MinD but with a varying diffusion coefficient for monomers (1.2 µm$^2$ s$^{-1}$), dimers (0.85 µm$^2$ s$^{-1}$), trimers (0.54 µm$^2$ s$^{-1}$), tetramers (0.34 µm$^2$ s$^{-1}$) and 0.3 µm$^2$ s$^{-1}$ for both penta- and hexamers. Distributions are grouped by localised particle density as a better estimate for the membrane crowdedness in a corresponding experimental video section. We used these oligomer-specific mass distribution shapes for the multicomponent fits in Fig. 2e-h and Supplementary Fig. 11. MinD and MinD D40A monomer detections are not possible at localised particle densities beyond 0.8 µm$^{-2}$. For the number of analysed trajectories, please refer to Supplementary Table 6.

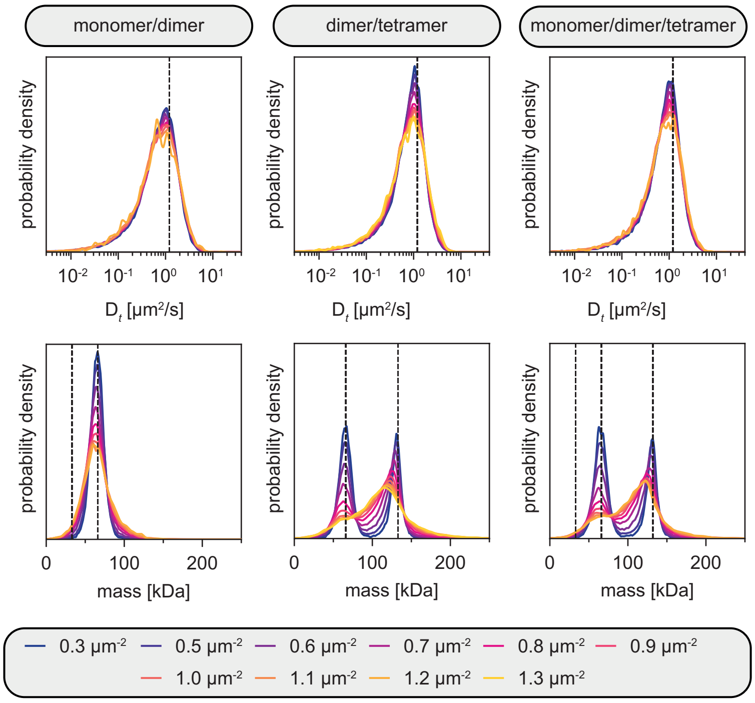


**Supplementary Figure 10 – Effect of mixtures of MinD oligomers as observed with simulated MSPT videos.** Distributions of determined diffusion coefficients (top row) and the extracted mass (bottom row) for videos containing simulated particles corresponding to mixtures of MinD oligomers. The simulation was set up with particles that had contrasts corresponding to MinD monomers and dimers (33 kDa – monomer, 66 kDa – dimer; left column) in a 1:1 mixture, a 1:1 mixture of MinD dimers and tetramers (132 kDa – tetramer, middle column), and a 1:1:1 composition of particles with the contrasts of a MinD monomer, dimer and tetramer (right column). Distributions are grouped by localised particle density. For the number of analysed trajectories, please refer to Supplementary Table 7.

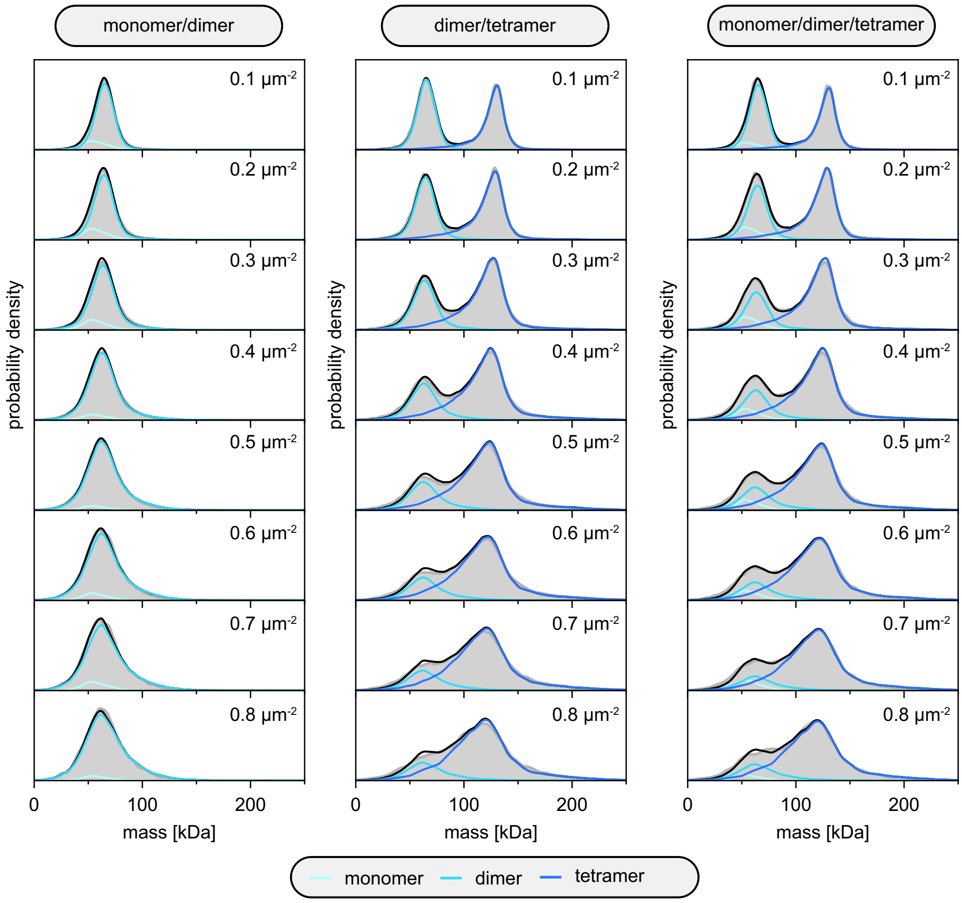


**Supplementary Figure 11 – Mass distributions of simulated particle mixtures fitted using separate oligomer components in Supplementary Fig. 9.** For each of the three simulated particle mixtures - monomer/dimer, dimer/tetramer, monomer/dimer/tetramer – the representative mass distributions (grey) and corresponding fit (black line, coloured lines highlight underlying components) is shown using a linear combination of the underlying separate components for eight different particle densities. For the number of analysed trajectories, please refer to Supplementary Table 7.

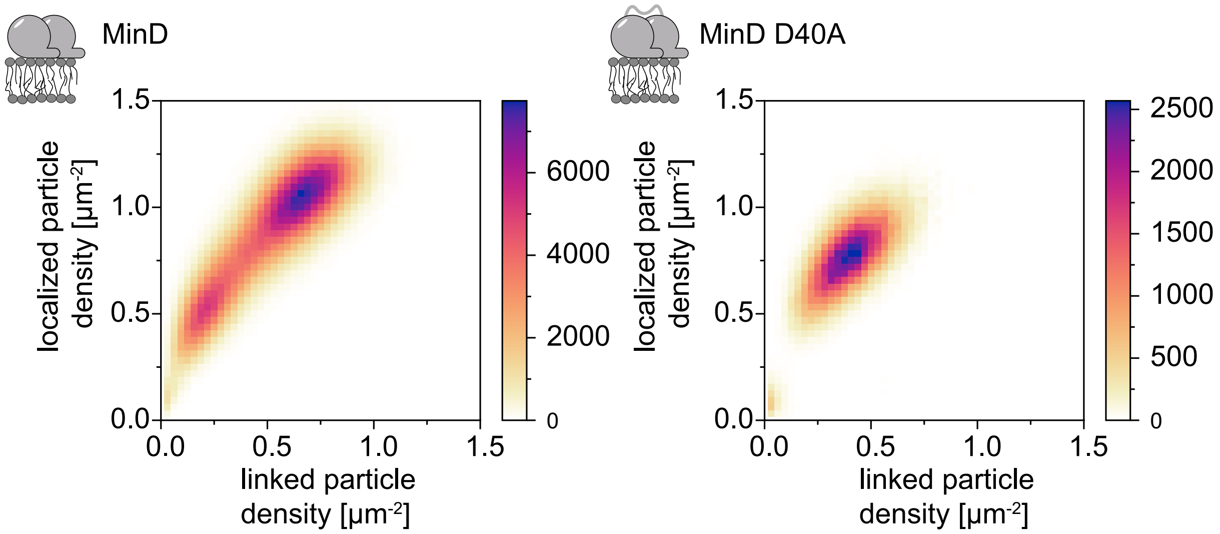


**Supplementary Figure 12 – Correlation of localised and linked particle densities for MinD and MinD D40A.** Comparison of frame-wise particle localisations per FOV to the linked particle density returned by the MSPT analysis. Note that linked particle densities are reduced by about 40% due to the custom-set trajectory length threshold of five frames and imperfect linking. Linked particle numbers: MinD n = 1,102,940 and MinD D40A n = 194,545,

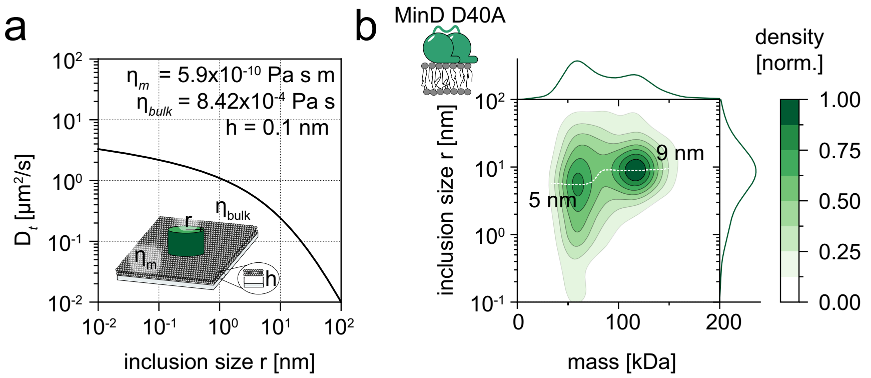


**Supplementary Figure 13 - MinD D40A tetramers insert all their subunit MTSs into the bilayer. (a)** Theoretical relationship between diffusion coefficient and inclusion size radius of an object in a supported lipid bilayer according to the theory of Evans-Sackmann [1], assuming a similar membrane viscosity ($\eta_m$) as for a pure DOPC membrane [2]. We have assumed $8.42 \times 10^{-4}$ Pa s [2] for the bulk viscosity $\eta_{bulk}$, and a height of 0.1 nm for the lubricating layer parameter h according to [1]. **(b)** Analysis of the inclusion size radius as a function of MinD D40A particle mass at a particle density of 0.1 $\mu m^{-2}$ (n = 12,063 trajectories). The dimer population had an estimated inclusion size radius of 5 nm, whereas the radius of the tetramer was 9 nm, indicating that the tetramer inserts additional MTSs into the bilayer. The white dashed line highlights the shift in inclusion size for increasing particle mass. Marginal probability distributions of both molecular mass and inclusion size radius are presented at the top and right, respectively.

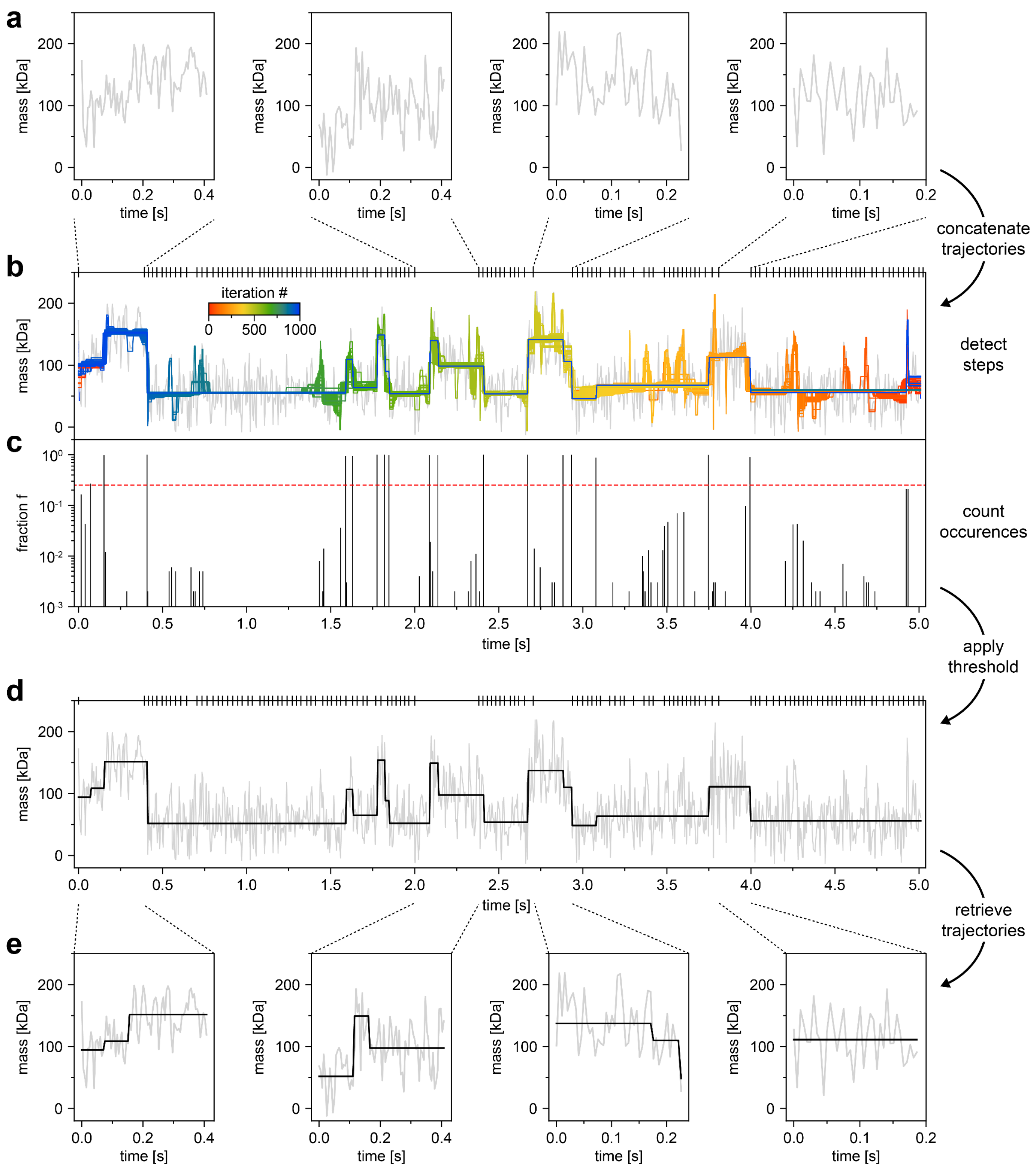


**Supplementary Figure 14 – Step detection workflow. (a)** Trajectories were concatenated in random order **(b),** beginning and end of individual trajectories is indicated by vertical lines at the top. Steps in subsections with a length of 1000 frames were identified using the Kalafut-Visscher algorithm[3], shifting the start point by one increment in each iteration. The significance of a step is resembled by its frequency of detection **(c)**, displayed as fraction *f* (frequency/1000) in logarithmic scale. Only steps that were detected more frequently than a threshold fraction (0.25) are retained **(d)**. The times series and its resulting step fit was then separated back into the original trajectories **(e)**.

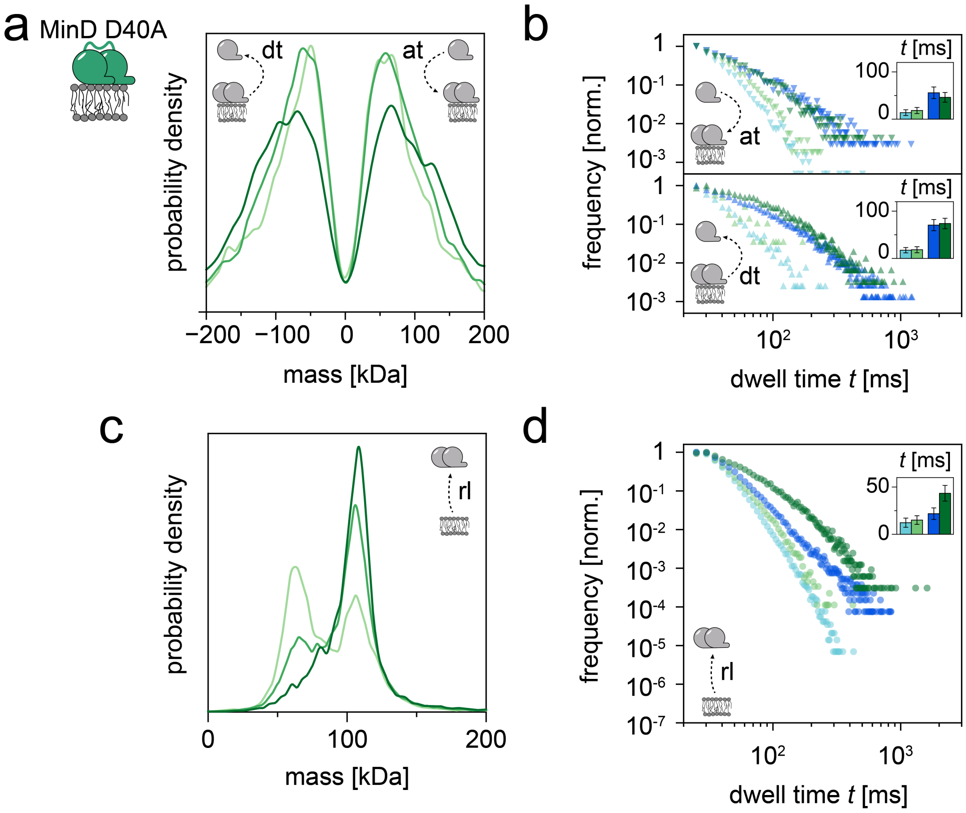


**Supplementary Figure 15 – Subunit (dis-)assembly of MinD D40A particles diffusing on a membrane. (a)** Mass step size distribution revealing MinD D40A subunit turnover at particle densities of 0.1 μm$^{-2}$ – pale green (n = 4,373 plateaus), 0.3 μm$^{-2}$ – light green (n = 17,443 plateaus), 0.5 μm$^{-2}$ – green (n = 4,746 plateaus). **(b)** Dwell time plots for MinD D40A and MinD subunit attachments (at) (top plot) and detachments (dt) (bottom plot). Dwell times are shown for the MinD dimer (light blue line) and MinD tetramer (dark blue line) as well as for their respective MinD D40A versions (dimer state – light green, tetramer state – green). Plateau numbers for (at): MinD dimer – n = 23,782, MinD D40A dimer – n = 6,862, MinD tetramer – n = 5,088, MinD D40A tetramer – n = 2,954; (dt): MinD dimer – n = 3,143, MinD D40A dimer – n = 295, MinD tetramer – n = 10,406, MinD D40A tetramer – n = 5,162. Insets: Mean dwell times of the two oligomer states. Error bars represent the standard deviation of the dwell times estimated by bootstrapping. **(c)** MinD D40A mass distribution for membrane release (rl) events at 0.1 μm$^{-2}$ – pale green (n = 12,063 plateaus), 0.3 μm$^{-2}$ – light green (n = 72,872 plateaus), 0.5 μm$^{-2}$ – green (n = 24,425 plateaus). **(d)** Plot of the dwell times before membrane release for the MinD D40A dimer and tetramer state and the respective MinD versions for comparison. Plateau numbers: MinD dimer – n = 562,011, MinD D40A dimer – n = 42,119, MinD tetramer – n = 73,037, MinD D49A tetramer – n = 28,467. Inset: Mean dwell times of the two oligomer states. Error bars represent the standard deviation of the dwell times estimated by bootstrapping.

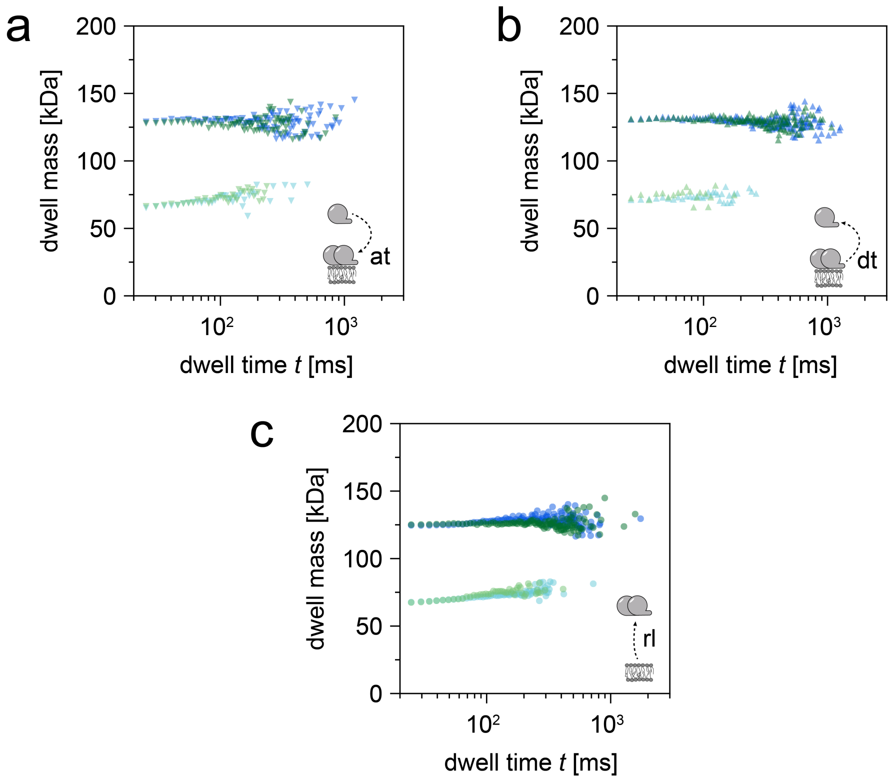


**Supplementary Figure 16 - Plots of dwell mass in relation to the dwell time for attachment events (a), detachment events (b) and membrane release (c), to illustrate mass-based categorisation of MinD D40A and MinD particles into dimer and tetramer states.** Dwell masses are shown for the MinD D40A dimer (light green) and MinD D40A tetramer (dark green) and for their respective wildtype versions MinD dimer (light blue line) and MinD tetramer (dark blue line). **(a)** Plateau numbers for attachment events (at): MinD dimer – n = 23,782, MinD D40A dimer – n = 6,862, MinD tetramer – n = 5,088, MinD D40A tetramer – n = 2,954 **(b)** Plateau numbers for detachment events (dt): MinD dimer – n = 3,143, MinD D40A dimer – n = 295, MinD tetramer – n = 10,406, MinD D40A tetramer – n = 5,162. **(c)** Plateau numbers for release events (rl): MinD dimer – n = 562,011, MinD D40A dimer – n = 42,119, MinD tetramer – n = 73,037, MinD D49A tetramer – n = 28,467.

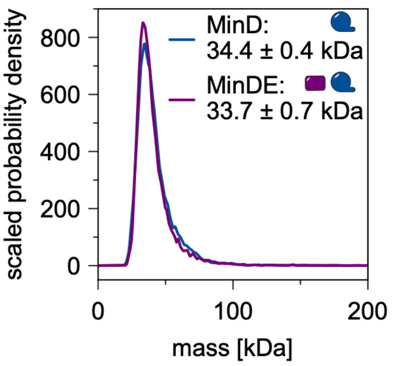


**Supplementary Figure 17 - In the absence of the catalytic membrane interface, MinD and MinE do not interact.** Probability density scaled by the number of detected molecules in videos of the same length for the mass distribution of 175 nM MinD (blue line; n = 16,013 particles), 175 nM MinDE (magenta line; n = 15,399) in the presence of 0.5 mM ATP.

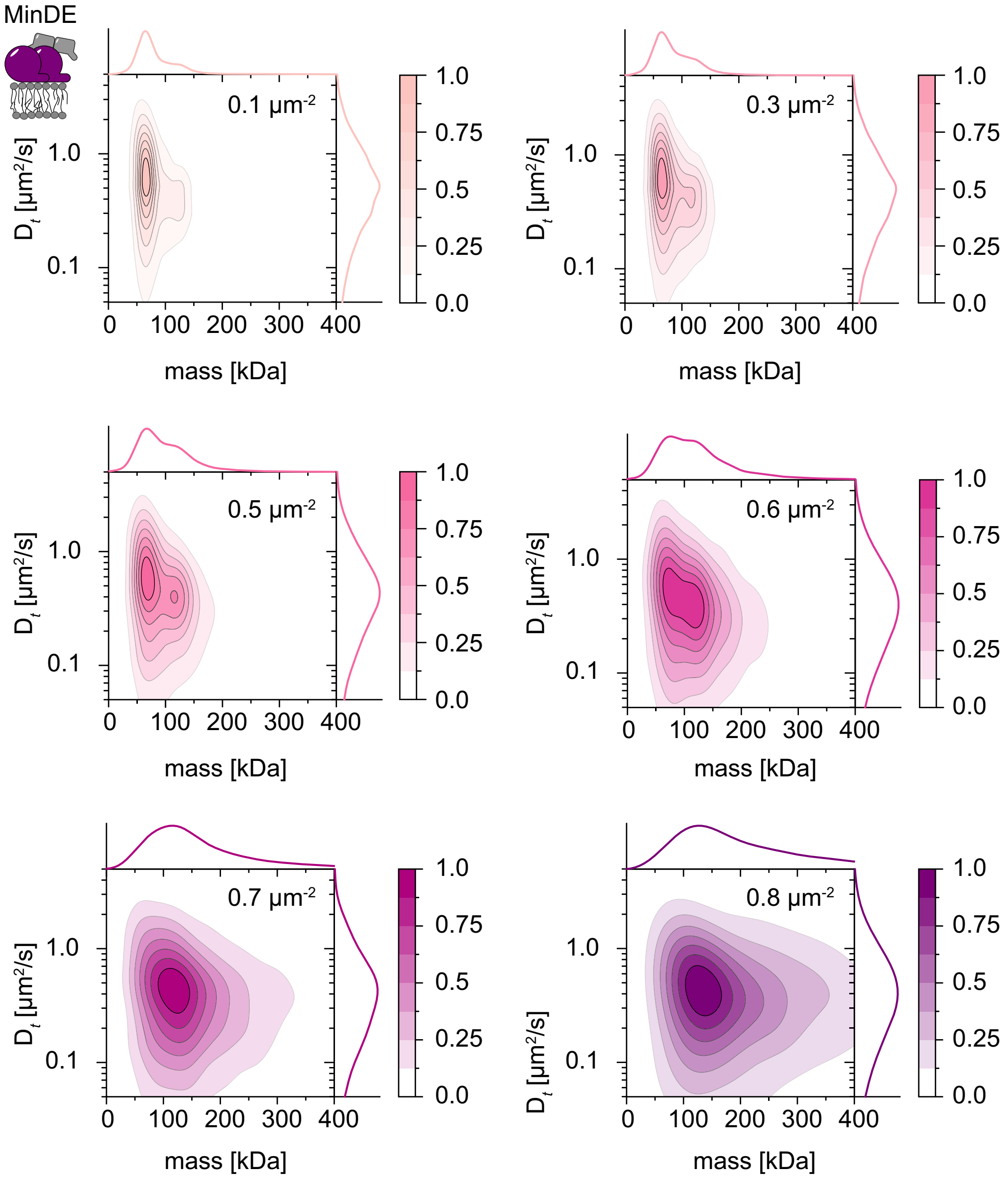


**Supplementary Figure 18 - 2D maps of diffusion coefficient and mass for membrane-bound MinDE particles.** 2D kernel density estimation of diffusion coefficient versus mass for membrane-attached MinDE at particle densities of 0.1 µm$^{-2}$ (light mauve, n = 200,436 trajectories), 0.3 µm$^{-2}$ (mauve, n = 158,660 trajectories), 0.5 µm$^{-2}$ (light pink, n = 35,501 trajectories), 0.6 µm$^{-2}$ (pink, n = 35,885 trajectories), 0.7 µm$^{-2}$ (light purple, n = 36,293 trajectories) and 0.8 µm$^{-2}$ (purple, n = 30,083 trajectories). Marginal probability distributions of both molecular mass and diffusion coefficient are presented at the top and right, respectively.

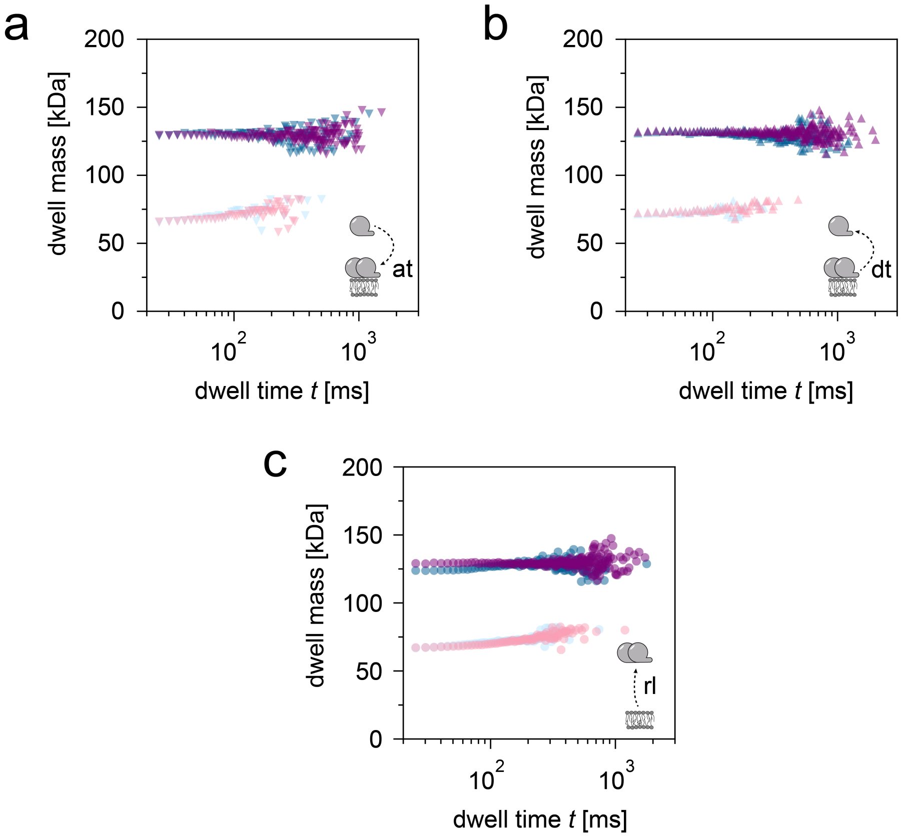


**Supplementary Figure 19 - Plots of dwell mass in relation to the dwell time for attachment events (a), detachment events (b) and membrane release (c), to illustrate mass-based categorisation of MinDE (pink shades) and MinD (blue shades) particles into dimer and tetramer states.** Dwell masses are shown for the MinD dimer (light blue line) and MinD tetramer (dark blue line) as well as for their respective MinDE complex versions (dimer state – light pink, tetramer state – pink). **(a)** Plateau numbers for attachment events (at): MinD dimer – n = 23,782, MinDE dimer – n = 37,278 and MinD tetramer – n = 5,088, MinDE tetramer – n = 11,698. **(b)** Plateau numbers for detachment events (dt): MinD dimer – n = 3,143, MinDE dimer – n = 3,974 and MinD tetramer – n = 10,406, MinDE tetramer – n = 22,501. **(c)** Plateau numbers for release events (rl): MinD dimer – n = 562,011, MinDE dimer – n = 277,782 and MinD tetramer – n = 73,037, MinDE tetramer – n = 60,114.

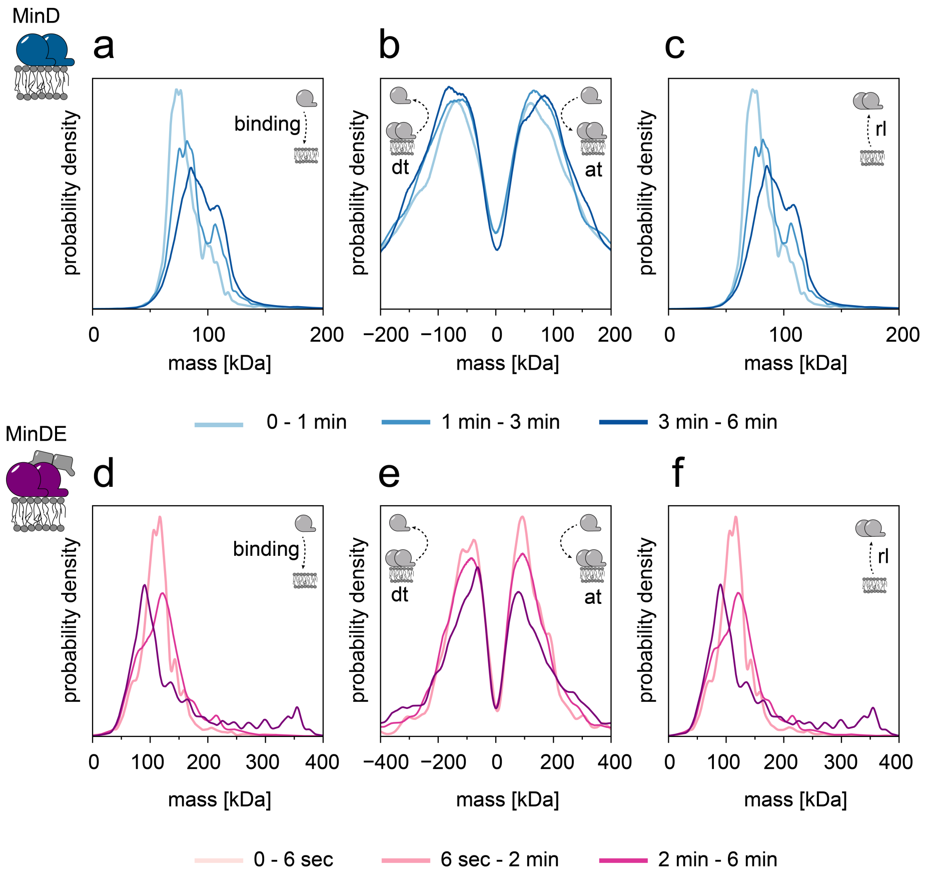


**Supplementary Figure 20 – Time-dependence of the membrane-associated behaviour of MinD and MinDE.** Plots depict the evolution of MinD (blue shades) and MinDE (purple shades) mass distributions for different time segments of a recorded video at a particle density of 0.5 µm$^{-2}$. MinD time segments: light blue – 0 to 1 min; blue – 1 to 3 min; dark blue – 3 to 6 min. MinDE time segments: light purple – 0 to 6 sec; purple – 6 sec to 2 min; dark purple – 2 min to 6 min. **(a, d)** MinD and MinDE mass distribution for membrane binding events. Plateau numbers for MinD (from light to dark blue): 53,176; 92,227; 139,510 and for MinDE (from light to dark purple): 5,161; 17,805; 12,535. **(b, e)** Mass step size distribution revealing the time-resolved subunit turnover of MinD and MinDE particles. Number of events for MinD (from light to dark blue): 4,630; 8,324; 12,910 and for MinDE (from light to dark purple): 1,087; 4,926; 3,234. **(c, f)** MinD and MinDE mass distribution for membrane release (rl) events. Number of plateaus for MinD (from light to dark blue): 53,176; 92,227; 139,510 and for MinDE (from light to dark purple): 5,161; 17,805; 12,535.

### Supplementary Discussion

For readers interested in applying the MSPT routine to their own iSCAT movies, we recommend to consider the details outlined in this Supplementary Discussion. As described in the methods section “Simulated videos of diffusing particles”, we set up a simulation that generates artificial videos of diffusing particles on an experimental SLB background. This simulation enabled us to test the performance of our MSPT analysis routine for different membrane scenarios.

We first assessed how well our analysis pipeline could reproduce the diffusion coefficient of particles with a size corresponding to MinD tetramers undergoing Brownian motion. By visual inspection of the videos, a median window size of 121 frames seemed sufficient to visualize particles diffusing at around 1 $\mu m^2 s^{-1}$, which was on the order of magnitude expected in our experimental videos. As pointed out earlier, the median-based background estimation works only if particles do not dwell for too long in a certain area. Therefore, we expected that the method would fail for lower diffusion coefficients. In fact, MSPT reproduced the input diffusion coefficients well between 0.32 $\mu m^2 s^{-1}$ and 10 $\mu m^2 s^{-1}$. However, below 0.32 $\mu m^2 s^{-1}$, coefficients were systematically overestimated (Supplementary Figure 2, left box, top row). The reason for this phenomenon becomes apparent when considering the extracted mass distributions (Supplementary Fig. 2, left box, bottom row). Since local background estimates get increasingly biased towards the value of background plus particle, if particles remain locally confined for the majority of the median period, the particles loose contrast at low diffusion coefficients. Consequently, masses were underestimated by the MSPT routine at diffusion coefficients below 0.32 $\mu m^2 s^{-1}$ and gradually fell below the detection level. Thus, the overestimation of slow diffusion coefficients is probably caused by fast particles with a higher SNR being more likely detected than the comparably slower ones with reduced SNR in the ensemble of simulated particle speeds.

The problem of particle confinement for the background estimation can be partially overcome by expanding the size of the median window. As shown in Supplementary Fig. 2, right box, both diffusion coefficient and particle mass were correctly extracted down to a diffusion speed of 0.1 $\mu m^2 s^{-1}$, if a median size of 1001 frames was chosen. Therefore, we recommend to carefully check that the median window size is chosen appropriately for the speed range of the imaged particles. Note, however, that large median windows drastically affect the image processing speed of the analysis routine and may become sensitive to sample drift.

We also compared the performance of two of the most common methods to extract diffusion coefficients from single-particle trajectories, namely MSD and JDD analysis (Supplementary Figure 2). We found that MSD performed slightly better, when extracting diffusion coefficients of slowly diffusing particles. However, for our experimental data with faster particles but many short trajectories we found that JDD produced narrower diffusion coefficient distributions. Therefore, we decided to employ this analysis method throughout the rest of the study.

In another scenario, we investigated the consequences of a crowded bilayer for our MSPT analysis. Similar to the confined particle scenario, we expected that the median-based background estimation would be affected, if many particles on the bilayer increased the probability that a location remains occupied by diffusing particles with diffusion coefficients between 0.3 to 1.2 $\mu m^2 s^{-1}$. Again, we compared the outcome for the distributions of diffusion coefficient and mass with the simulation inputs (Supplementary Figs. 9 and 10). While the estimated diffusion coefficients remained correct, we observed an increasing distortion of the mass distributions towards low masses at high particle densities. This effect was most pronounced for particles with high contrast. Dense bilayers probably lead to a similar situation as with particle confinement, where the median-based background estimate locally approaches the value of background plus particle, thus reducing apparent particle contrasts in the ratiometric movie. For this reason, we recommend to perform MSPT experiments at localised particle densities ideally below 0.7 $\mu m^{-2}$.

As we were interested in quantifying the oligomer distribution of MinD at protein densities beyond this limit, we could not assume a simple Gaussian mixture model underlying the mass distributions in Figures 2e/g, considering the observed distribution shape changes in our simulation. Fortunately, we found in a simulation with mixtures of particles with different contrasts that their respective peaks kept a particle-density-specific shape that was similar to the shape for the individual species at that particle density (Supplementary Fig. 10). Accordingly, we could use the simulated numerical form of a species mass distribution at a given particle density to fit a linear combination of these distributions with varying amplitudes to describe the mass distribution of a species mixture and extract their relative abundances (see Supplementary Fig. 11 for a fit of simulated data and Fig. 2e-h for a fit of our experimental data).

While this approach enables the deconvolution of complex mass distributions such as those observed for MinD at higher particle densities, where the underlying populations cannot be resolved experimentally, the remaining caveat is that this approach does not improve the particle detection efficiency itself. Hence, the abundance of small protein complexes close to the detection limit such as MinD dimers tends to be underestimated at high particle densities (Supplementary Figures 10 and 11).

### Supplementary Tables

**Supplementary Table 1** Particle detection and trajectory linking parameters for MSPT measurements.

| Particle detection parameters | | Trajectory linking parameters (Trackpy link_df function) | |
|---|---|---|---|
| median half-size n | 60 | search_range | 4 pixels |
| Laplace filter threshold | 0.0007 | memory | 0 frames |
| Laplace filter sigma | 1.5 pixels | | |
| Local maximum filter size | 7 pixels | | |
| Refeyn PSF parameters | A12: -5.9360386376<br>W = 2.1436230397<br>S = 12.8930119585 | | |

**Supplementary Table 2** List of the molecular mass and the determined iSCAT contrast of standard proteins measured using the conventional MP landing assay.

| standard protein | used abbreviation | MW [kDa] | iSCAT contrast |
|---|---|---|---|
| alcohol dehydrogenase | ADH | 147.4 | 0.0053 ± 0.0002 |
| bovine serum albumin | BSA | 66.4 | 0.0025 ± 0.0002 |
| TEV protease | TEV | 28.6 | 0.0015 ± 0.0002 |
| β-amylase | bAm | 224.3 | 0.0080 ± 0.0010 |
| protein A | prA | 42.0 | 0.0019 ± 0.0002 |

**Supplementary Table 3** List of the molecular mass and the determined iSCAT contrast of standard proteins measured using MSPT.

| standard protein | used abbreviation | expected MW [kDa] | iSCAT mass | iSCAT contrast |
|---|---|---|---|---|
| divalent streptavidin | Strep | 55.2 | 52.2 ± 8.8 | 0.0022 ± 0.0004 |
| divalent streptavidin with biotinylated aldolase | Strep-ALD | 133.9 (2x ALD)<br>212.5 (4x ALD) | 131.7 ± 13.0<br>193.0 ± 26.4 | 0.0049 ± 0.0005<br>0.0069 ± 0.0009 |
| divalent streptavidin with biotinylated bovine serum albumin | Strep-BSA | 124.5 (1x BSA)<br>193.8 (2x BSA) | 127.0 ± 15.3<br>204.5 ± 22.2 | 0.0047 ± 0.0006<br>0.0072 ± 0.0008 |
| divalent streptavidin with biotinylated protein A | Strep-prA | 101.9 | 111.9 ± 15.9 | 0.0042 ± 0.0006 |
| tetravalent streptavidin | - | 55.2 | 55.1 ± 1.2 | 0.0023 ± 0.0001 |

**Supplementary Table 4** List of all plasmids that were used for the characterisation of the MinDE mechanism through MSPT.

| vector | protein construct | source |
|---|---|---|
| pET28a-MinE-His | MinE-His | [4] |
| pET28a-His-MinD-MinE | MinD | [5] |
| pET28a-MinD(D40A)-MinE | MinD(D40A) | [6] |
| pET21a-Streptavidin-Alive | Streptavidin-Alive | [7]; Addgene plasmid #20860 |
| pET21a-Streptavidin-Dead | Streptavidin-Dead | [7]; Addgene plasmid #20859 |

**Supplementary Table 5** Number of detected trajectories, of simulated particles with varying diffusion coefficients. Videos of simulated particles were either analysed with a median window of 121 or 1001 frames.

| | Median window of 121 frames | | Median window of 1001 frames | |
|---|---|---|---|---|
| $D_t$ [µm²/s] | N(trajectories) | | N(trajectories) | |
| 0.01 | 4,143 | 2,210 | 5,415 | 3,891 |
| 0.032 | 14,525 | 10,401 | 3,692 | 3,004 |
| 0.1 | 10,072 | 8,492 | 4,470 | 3,781 |
| 0.32 | 10,498 | 9,273 | 6,739 | 5,910 |
| 1.0 | 17,090 | 15,324 | 10,637 | 9,558 |
| 3.2 | 20,881 | 19,071 | 13,554 | 12,368 |
| 10.0 | 28,880 | 25,613 | 19,030 | 16,816 |

**Supplementary Table 6** Number of detected trajectories, of simulated MinD and MinD D40A particles in respect to the particle density.

| | monomer | dimer | trimer | tetramer | pentamer | hexamer |
|---|---|---|---|---|---|---|
| particle density [µm²] | N(trajectories) | | | | | |
| **MinD** | | | | | | |
| 0.3 | 11,524 | 17,892 | 19,604 | 18,958 | 19,402 | 19,654 |
| 0.5 | 26,531 | 52,722 | 55,829 | 49,248 | 57,528 | 57,868 |
| 0.6 | 15,895 | 59,883 | 65,019 | 52,919 | 66,897 | 66,872 |
| 0.7 | 4,752 | 73,411 | 63,690 | 60,632 | 62,524 | 65,148 |
| 0.8 | - | 103,375 | 66,002 | 73,141 | 63,726 | 64,981 |
| 0.9 | - | 144,803 | 84,756 | 82,659 | 83,495 | 81,390 |
| 1.0 | - | 159,423 | 101,707 | 74,458 | 101,969 | 102,317 |
| 1.1 | - | 123,062 | 99,319 | 49,996 | 99,241 | 102,678 |
| 1.2 | - | 59,992 | 72,792 | 22,537 | 71,813 | 75,321 |
| 1.3 | - | 18,013 | 37,098 | 6,119 | 34,951 | 36,882 |
| **MinD D40A** | | | | | | |
| 0.3 | 11,524 | 18,938 | 5,905 | 17,100 | 5,585 | 6,014 |
| 0.5 | 26,531 | 59,486 | 16,059 | 49,522 | 14,765 | 15,505 |
| 0.6 | 15,895 | 74,810 | 15,600 | 52,183 | 14,632 | 15,552 |
| 0.7 | 4,752 | 75,636 | 14,442 | 50,917 | 15,464 | 16,591 |
| 0.8 | - | 79,327 | 19,142 | 62,124 | 19,170 | 18,976 |
| 0.9 | - | 95,539 | 24,806 | 76,452 | 22,748 | 21,562 |
| 1.0 | - | 101,936 | 27,228 | 85,122 | 22,984 | 22,374 |
| 1.1 | - | 79,968 | 25,654 | 78,347 | 20,450 | 21,985 |
| 1.2 | - | 41,960 | 18,553 | 52,117 | 14,528 | 16,721 |
| 1.3 | - | 14,680 | 9,745 | 26,031 | 6,868 | 8,480 |

**Supplementary Table 7** Number of detected trajectories of simulated particles in respect to the particle density. The simulation was set up with particles that had theoretical masses corresponding to a 1:1 mixture of MinD monomers and dimers, a 1:1 mixture of MinD dimer and tetramer, and a 1:1:1 composition of particles with the MW of a MinD monomer, dimer and tetramer.

| | **monomer/dimer** | **dimer/tetramer** | **monomer/dimer/tetramer** |
|---|---|---|---|
| **Particle density [$\mu m^2$]** | **N(trajectories)** | | |
| 0.1 | 81,389 | 83,453 | 71,449 |
| 0.2 | 84,518 | 87,634 | 68,765 |
| 0.3 | 92,679 | 89,402 | 81,044 |
| 0.4 | 94,099 | 105,408 | 92,079 |
| 0.5 | 71,475 | 112,960 | 77,079 |
| 0.6 | 34,918 | 100,613 | 44,770 |
| 0.7 | 9,714 | 62,044 | 16,675 |
| 0.8 | 1,738 | 24,813 | 3,722 |

**Supplementary Movies**

**Supplementary Movie 1 -** Comparison of processed movies showing a single diffusing particle of biotin-aldolase bound to a biotinylated bilayer via divalent streptavidin. Left: processing strategy typically used for mass photometry (sliding mean, navg = 5, Supplementary Fig. 1a and b). Right: new processing strategy using a sliding median as background estimate (median half-size = 60 frames, Supplementary Fig. 1c). Scale bar: 1 μm. Interferometric scattering contrast range: black = 0.01; white = −0.008.

**Supplementary Movie 2 -** Exemplary movies showing standard proteins diffusing on a biotinylated bilayer attached via divalent streptavidin. Upper left: Divalent streptavidin alone. Upper right: biotin-BSA-streptavidin. Lower left: biotin-protein A-streptavidin. Lower right: biotin-aldolase-streptavidin. Image processing median half-size = 60 frames. Scale bar: 1 μm. Interferometric scattering contrast range: black = 0.004; white = −0.003.

**Supplementary Movie 3 -** Exemplary movies showing MinD (top) and MinDE (bottom) complexes diffusing on a bilayer. Solution concentrations: top - 100 nM MinD; bottom - 100 nM MinD, 100 nM MinE. Image processing median half-size = 60 frames. Scale bar: 1 μm. Interferometric scattering contrast range: black = 0.004; white = −0.003.

### References


1. Evans, E. & Sackmann, E. Translational and rotational drag coefficients for a disk moving in a liquid membrane associated with a rigid substrate. *J. Fluid Mech.* **194**, 553–561 (1988).
2. Khmelinskaia, A., Mücksch, J., Petrov, E. P., Franquelim, H. G. & Schwille, P. Control of Membrane Binding and Diffusion of Cholesteryl-Modified DNA Origami Nanostructures by DNA Spacers. *Langmuir* **34**, 14921–14931 (2018).
3. Little, M. A. *et al.* Steps and bumps: Precision extraction of discrete states of molecular machines. *Biophys. J.* **101**, 477–485 (2011).
4. Glock, P. *et al.* Stationary patterns in a two-protein reaction-diffusion system. *ACS Synth. Biol.* **8**, 148–157 (2019).
5. Loose, M., Fischer-Friedrich, E., Ries, J., Kruse, K. & Schwille, P. Spatial regulators for bacterial cell division self-organize into surface waves in vitro. *Science* **320**, 789–792 (2008).
6. Heermann, T., Ramm, B., Glaser, S. & Schwille, P. Local Self-Enhancement of MinD Membrane Binding in Min Protein Pattern Formation. *J. Mol. Biol.* **432**, 3191–3204 (2020).
7. Howarth, M. *et al.* A monovalent streptavidin with a single femtomolar biotin binding site. *Nat. Methods* **3**, 267–273 (2006).
